# Spore morphology and evolution in *Isoetes* (Isoetales)

**DOI:** 10.1101/2024.12.31.630945

**Authors:** Eva Larsén, Anbar Khodabandeh, Catarina Rydin

## Abstract

*Isoetes* (Isoetaceae, Isoetales) is a cosmopolitan genus with an ancient and diverse evolutionary history that presumably peaked in the Paleozoic. The age of the living clade has never been satisfactorily clarified. With little morphological (and molecular) divergence among species, megaspore morphology may provide valuable information on relationships in the group, not least among extant and extinct forms. We study megaspore ornamentation and surface texture/structure in 74 phylogenetically placed samples representing 59 species of *Isoetes* using SEM, and we discuss evolutionary implications of the results. Ornamentation (classified into 12 categories) and surface structure/texture (10 categories) were mapped onto a phylogeny constructed based on molecular data from the same samples whenever possible. All megaspores of the family are trilete with an outermost siliceous coating. There is ample micromorphological variation, of which some appears clade specific. The megaspores of *Isoetes reticulata*, discovered from the late Oligocene to early Miocene of Tasmania, share remarkable similarities with those of extant species in the Australasian clade, in particular *Isoetes neoguineensis* and partly also *Isoetes japonica*. We argue that this fossil could be used to calibrate the age of the Australasian crown group (clade D) to absolute time (c. 20-25 Ma) in analyses of divergence times of clades based on molecular data.

## INTRODUCTION

*Isoetes* L. (Isoetaceae, Isoetales) is a cosmopolitan genus of heterosporous lycopods with a most commonly semi-aquatic lifestyle (e.g. Pfeiffer 1922, Taylor and Hickey 1992, Rydin and Wikström 2002, Hoot *et al*. 2004, Troìa *et al*. 2016). The genus belongs to the order Isoetales, which appeared in the Late Devonian and had its greatest diversity and ecological importance in the late Carboniferous when arborescent lepidodendrids dominated the coal swamps of Euramerica (e.g. Bateman *et al*. 1992, Pigg 2001). Living alongside and outliving their tree-shaped relatives were also several genera of smaller and mostly unbranched isoetaleans that were widespread in the Triassic and continued to diversify throughout the Mesozoic and Cenozoic, up to the present day (e.g. Bateman *et al*. 1992, Pigg 2001).

The extant genus *Isoetes* shares several synapomorphies with their fossil relatives such as pseudobipolar growth from a shoot-like rootstock, stigmarian root systems with dichotomizing roots and leaflike lateral rootlets borne in helical rhizotaxy, and secondary xylem produced by a unifacial cambium (e.g. Bateman *et al*. 1992, DiMichele and Bateman 1996, Kenrick and Crane 1997, DiMichele *et al*. 2022). The c. 200 species (e.g. Larsén and Rydin 2016, PPG 1 2016, Troìa *et al*. 2016) of the genus can vary greatly in size but morphological divergence (as well as molecular sequence divergence) is otherwise limited (e.g. Rydin and Wikström 2002, Hoot *et al*. 2004, Larsén and Rydin 2016, Troìa *et al*. 2016). They are characterised by a short, woody corm that produce dichotomously branching roots and spirally arranged leaves (e.g. Pfeiffer 1922, Jermy 1990). The leaves are linear and usually contain four air channels longitudinally as well as a ligule above a basal and sunken sporangium. The sporangium can be covered by a velum and produces either many small microspores or fewer large megaspores (e.g. Pfeiffer 1922, Jermy 1990).

The species of *Isoetes* usually grow in damp to boggy habitats, presumably reflecting their ancestry in swamp forests of the Carboniferous but while most species inhabit seasonally inundated areas, there are terrestrial and fully aquatic species too (e.g. Pfeiffer 1922, Jermy 1990, Rydin and Wikström 2002). Habitat preferences were used for subgeneric classification by some authors (e.g. Baker 1880, Engelmann 1882, Motelay and Vendryès 1884), however, Duthie was early to criticise the delimitation of *Isoetes* species based on habitat (Duthie 1929). She argued that the ecological preferences used by these authors are not adequate for subgeneric divisions in *Isoetes*. Also in more recent work it has been discussed that there are several indications of a complex evolutionary history regarding habitat preferences of species of the genus (e.g. Taylor and Hickey 1992, Keeley *et al*. 1994, Rydin and Wikström 2002), probably involving multiple evolutionary transitions between habitats during the evolution of the group, as e.g. indicated by differences among species regarding stomata distributions and metabolism.

Evolutionary studies and species determination in *Isoetes* are difficult tasks since both morphological and molecular divergence among species is generally low (e.g. Rydin and Wikström 2002, Schuettpelz and Hoot 2006, Larsén and Rydin 2016, Pereira *et al*. 2021, Larsén *et al*. 2022). Among varying and potentially useful morphological characters are for example the corm and its lobes (Freund *et al*. 2018). This feature has been found to vary within populations of at least some species (Croft 1980) so its usefulness for evolutionary studies of *Isoetes* at the genus level could probably be questioned. Freund *et al*. (2018) found a phylogenetic diagnostic character in the number of lobes of young plants, but this was only useful for telling clade E of Larsén and Rydin (2016), i.e. the North and South American clade, apart from other clades. The length of the leaves is also sometimes used to tell species apart but again this feature can vary between populations and additionally the ranges can overlap (Croft 1980). As for megasporangium size and megaspore number per sporangium, these characteristics vary with the age of the plant (Budke *et al*. 2005) and for a perennial plant like *Isoetes* that only has secondary growth in the small corm, age can be tricky to tell.

Since it is hard to estimate the phylogenetic position of extant *Isoetes* species based on morphology it is an even bigger problem when considering fossil taxa as there is no available DNA to sequence for them. Ample work has been conducted to elucidate the evolutionary history of the isoetalean lineage (e.g. Hickey 1986a, Bateman *et al*. 1992, DiMichele and Bateman 1996, Kenrick and Crane 1997, DiMichele and Bateman 2020), but it has also been repeatedly pointed out that the Mesozoic and Cenozoic fossils of the group are less well understood and relationships among them, the Paleozoic fossils, and the contemporary species of *Isoetes* remain difficult to assess (e.g. Ash and Pigg 1991, Pigg 2001, Taylor *et al*. 2009, DiMichele and Bateman 2020), potentially including ample parallel evolution and difficulties to understand the homology of features.

In a genus like *Isoetes* with species that may be cryptic or at least hard to distinguish based on gross morphology, the megaspores are often used to ascertain the identification of the species (Proctor 1949). And megaspore morphology has indeed been used to create subgroups within *Isoetes*, e.g., by Pfeiffer (1922) in her monograph over the genus. Furthermore, megaspore morphology could constitute a possible link between living and fossil species in evolutionary studies if any phylogeny-wide characteristics could be found. Skog and Hill (1992) point out that fossil megaspores are often not preserved with the silicified coating that extant species commonly possess, potentially leading to misinterpretations of the homology of traits. However, Troìa *et al*. (2012) analysed Mediterranean *Isoetes* species’ megaspores and compared their untreated appearance versus that after treatment with acid to remove the siliceous outer coating, and found that the distinctive ornamentation was still visible underneath although much fainter.

Nevertheless, ample work remains before megaspore morphology can be effectively used in evolutionary studies of *Isoetes* and the isoetalean order. While Musselman (2002) made a comprehensive overview of microspore micromorphology and ultrastructure, no similar study as concerns phylogenetic scope has been undertaken on megaspore morphology since Pfeiffer’s classificational work (1922). Pfeiffer’s study (1922) is very impressive considering both the number of species included, spores described, and the level of detail. But it cannot be helped that without the availability of DNA sequencing techniques and scanning electron microscopy it is difficult to go beyond descriptive work, which in addition will have a lower resolution and level of details compared to more recent work. Since Pfeiffer (1922) there has been many studies that have improved our understanding of *Isoetes*’ megaspores (e.g. Marsden 1979, Croft 1980, Kott and Britton 1983, Hickey 1986b, Britton *et al*. 1999, Holmes *et al*. 2005, Troìa *et al*. 2012, Pereira and Labiak 2013, Brunton 2015, Pereira *et al*. 2016) and although these studies have added immeasurably to the knowledge of the genus the focus has been on smaller geographical regions and comparison of co-occurring species within those areas.

Here we study the megaspore morphology of phylogenetically placed *Isoetes* species (Larsén *et al*. 2022). We describe the ornamentation and surface texture/structure of the megaspore of 74 specimens (59 species), representing the vast majority of the global phylogenetical and geographical diversity of *Isoetes*. We elaborate on evolutionary patterns regarding these features across the genus tree, and we briefly discuss implications of our results for the understanding of the evolution of isoetaleans through time.

## MATERIALS AND METHODS

### Taxon sample and sampling strategy

Our sampling strategy was to examine the spores from specimens that were included in our previous phylogenetic studies based on molecular data (Larsén and Rydin 2016, Larsén *et al*. 2022) so that their phylogenetic position within the genus is known. Occasionally additional specimens were studied if the DNA-sampled specimen lacked developed megaspores or if they were in a too deteriorated state to permit sufficient investigations. Effort was made to make sure that the alternative spore sample was phylogenetically congruent with the DNA-sampled individual. In the case of *Isoetes malinverniana* Ces. & De Not., several specimens were sampled to ascertain that the appearance of an extra, raised layer of netted fibers on its spores is consistent, not simply a feature of a single sample. Investigated samples (taxon names, species distributions, DNA voucher information including area and year of collection, and lab identity numbers) are given in Supporting Information Table S1. Authors of names follow the recent checklist by Troìa *et al*. (2016); see also nomenclatural notes in Larsén *et al*. (2022).

### SEM studies

In order to avoid artificial alteration of the spores (Norbäck Ivarsson 2013, Bolinder *et al*. 2016) no pre-treatment of the spores was conducted. Air-dried spores from a single sporangium were placed on double-sided tape on aluminium stubs, sputter-coated with gold and examined with a Hitachi-TM3000 SEM (at the Electron Microscopy Center, Stockholm University). The ornamentation and surface texture/structure were documented and the equatorial diameter of the spores was measured during the SEM studies. The equatorial diameter is here defined as the distance from the tip of one arm of the proximal ridge to the opposing radial side, including the equatorial ridge. Fungal and bacterial growth was common, but easy to tell apart from the spore ornamentation. Some spores were clearly degraded by age (e.g., sample ELS93) but we rarely encountered this problem. In addition, we studied freshly collected spores of *Isoetes lacustris* L. and compared their morphology with those of older collections of the same species and found no relevant differences. The terminology used to describe the spores mainly follows Hickey (1986b). The most important general terms are illustrated in Figure 1.

**Figure 1.**
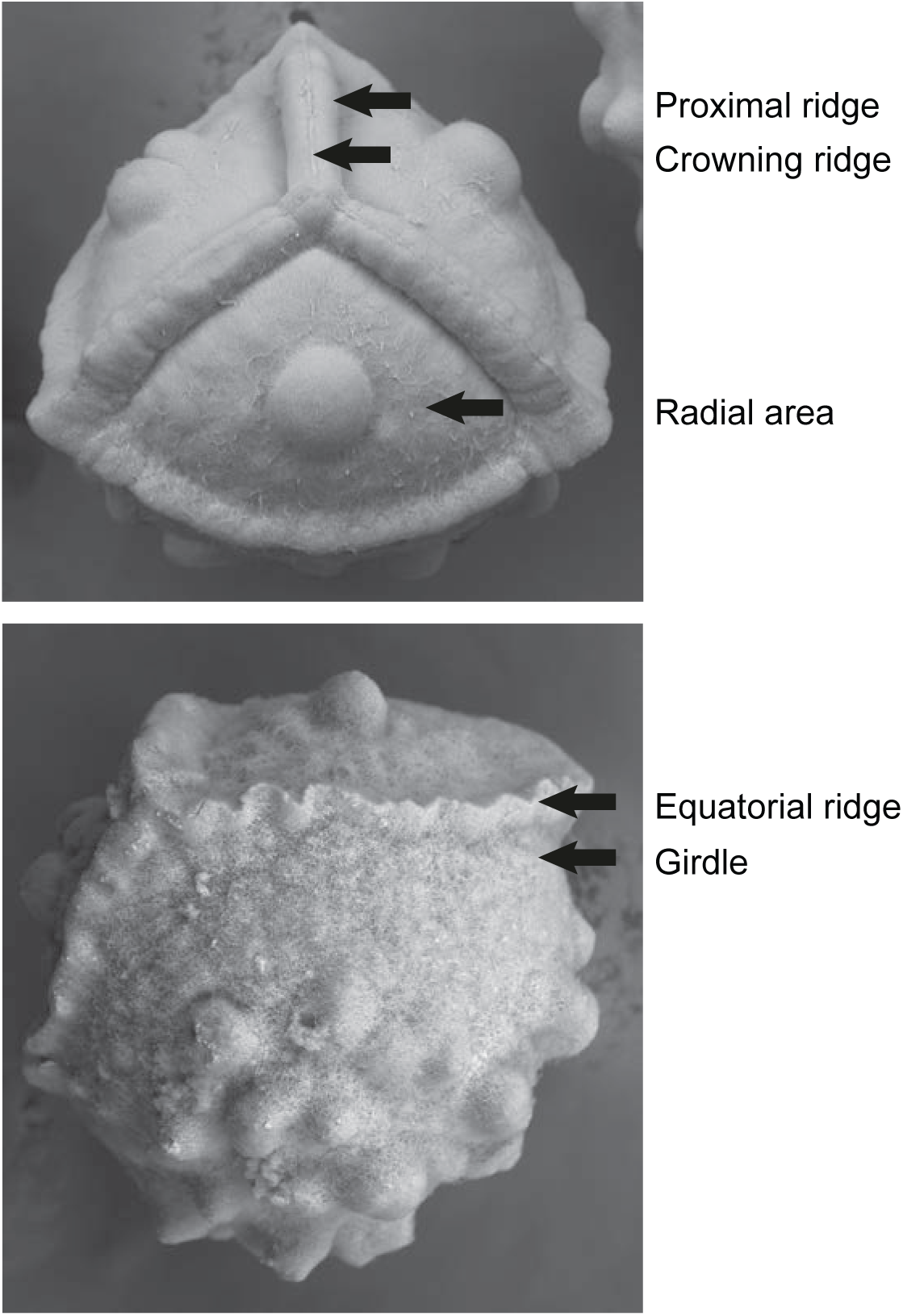
Key morphological structures and general terms used here to describe megaspore morphology in *Isoetes*. The terminology used mainly follows Hickey (1986b).

### Molecular data and analysis

Data from seven molecular markers (plastid *ndhC-ndhK*, *rbcL*, *rpoC1*, *ycf1*, *ycf66*, and *trnV*^UAC^ including its subsequent spacer, and the nuclear ribosomal internal transcribed spacer (nrITS)) were utilized to produce a phylogenetic framework for our study. Most of these sequences were produced and used by us in a previous study (Larsén *et al*. 2022), but new sequences of nrITS were produced here for a few samples (EL085, EL163, EL164, EL165, EL167 and EL168) as well as one sequence of *rpoC1* for EL114 and analysed together with sequences downloaded from GenBank (https://www.ncbi.nlm.nih.gov/genbank/). See Supporting Information Table S1 for voucher information and GenBank accessions of utilized data.

Extraction of total genomic DNA was performed according to the cetyltrimethyl-ammonium bromide (CTAB) method (Doyle and Doyle 1987, Doyle 1991) and purified using a QIAquick PCR Purification Kit (Qiagen, Sweden). PCR reactions were conducted using standard procedures (as described for example in Thureborn *et al*. 2019) and optimized for the here utilized primers and extractions. Sequencing was performed by the Macrogen Sequencing Service (Amsterdam). Obtained sequences were aligned by eye to the matrices produced by Larsén *et al*. (2022) in Geneious version 2022.2.2 (www.geneious.com).

ModelFinder (Kalyaanamoorthy *et al*. 2017) as implemented in the IQ-TREE web server (Trifinopoulos *et al*. 2016) was used to estimate the best fitting models and partitions (Chernomor *et al*. 2016). Best fitting substitution models (the criteria AIC, AICc and BIC gave similar scores) for the two edge-linked partitions nrITS and plastid regions were TPM3u+F+G4 and GTR+F+I+G4, respectively. Maximum likelihood analyses were conducted on the IQ-TREE web server (Trifinopoulos *et al*. 2016). Bootstrap support values were obtained using Ultrafast bootstrap (Hoang *et al*. 2018) as implemented in IQ-TREE 2 (Minh *et al*. 2020) with number of bootstrap alignments set to 1000, maximum likelihood iterations set to 1000, minimum correlation coefficient set to 0.99 and other settings at default values.

## RESULTS

### Megaspore descriptions

The megaspores of *Isoetes* are trilete, and have a distinct equatorial ridge, which in some cases is subtended by a girdle. The samples are described in the order of appearance on the phylogeny (Fig. 2). The overall ornamentation is based on observations of the distal side if the sides vary. Scanning electron micrographs of investigated samples are provided (Figs 3-21).

**Figure 2.**
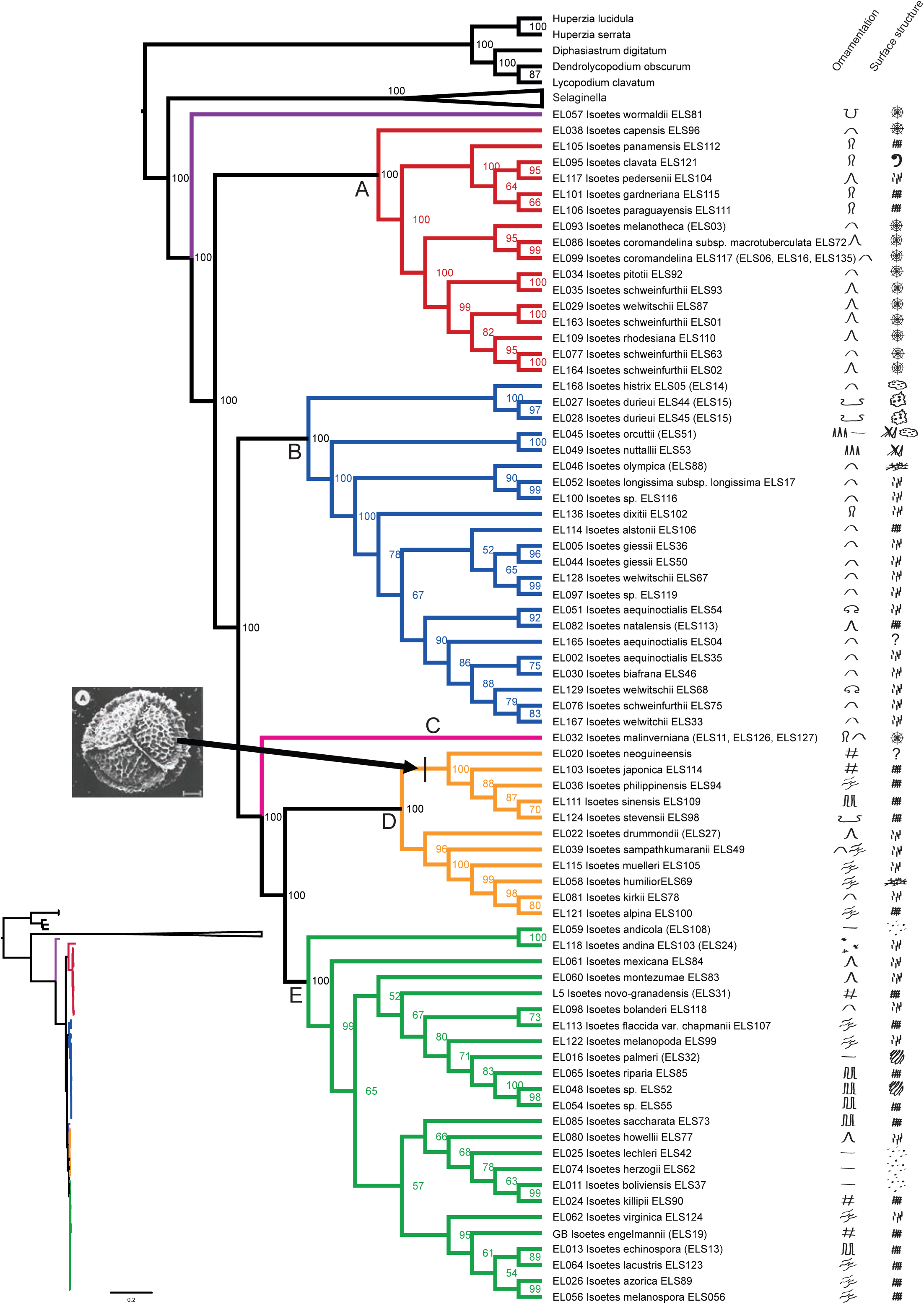
Phylogeny with megaspore ornamentation and surface structure indicated to the right. Phylogram inset to the lower left. DNA laboratory-id is shown before the species name and spore-id after (a spore-id in parenthesis indicates that another specimen than the sequenced one has been studied). Inset photo depicts a fossil spore of *Isoetes reticulata* (Hill 1988), reproduced with permission from the publisher. Its hypothesized phylogenetic position is indicated. Ornamentation symbols: 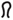=baculate; 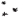=cristate; 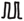=echinate; 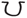=foveolate; 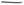=levigate; 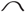=pustulate; 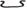=retate; 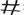=reticulate; 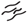=rugulate; 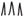=spiculate; 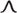=tuberculate; 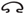=umbellate. Surface structures: 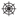=cobweb; 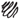=dense; 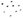= dense with pores; 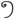=fibers forming curls; 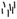=hairy; 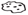=melted cheese; 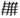=network; 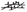=network with loose fibers; 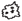= sponge-like; 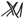=thorny.

**Figure 3.**
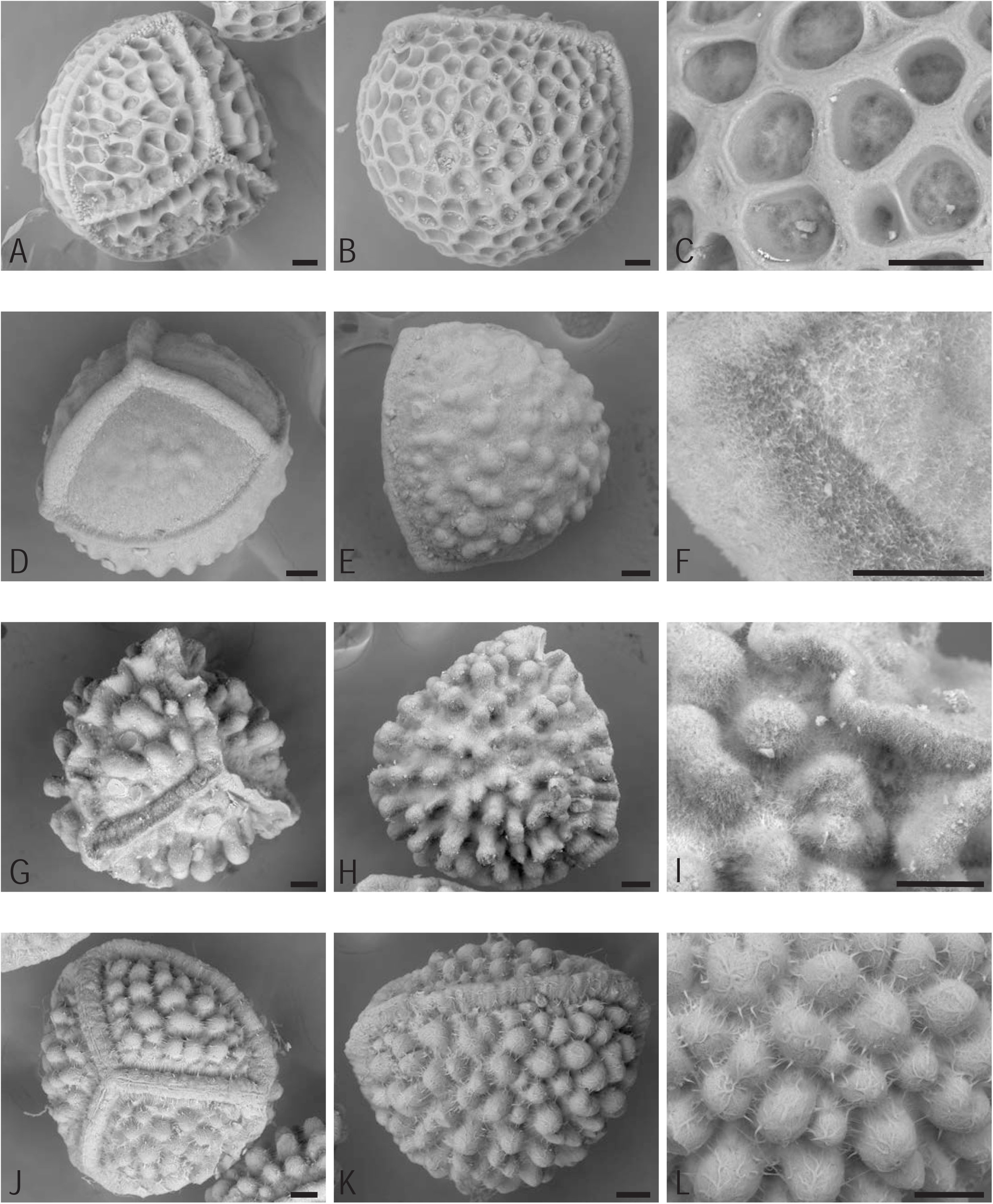
*Isoetes wormaldii* and species of clade A. All scale bars are 50 μm A-C *Isoetes wormaldii* ELS81, A proximal side, B distal side, C magnified ornamentation on distal side. D-F *Isoetes capensis* ELS96, D proximal side, E distal side, F magnified surface structure on proximal side. G-I *Isoetes panamensis* ELS112, G proximal side, H distal side, I magnified ornamentation on distal side. J-L *Isoetes clavata* ELS121, J proximal side, K distal side, L magnified ornamentation on distal side.

#### *Isoetes wormaldii* Sim ELS81 (Fig. 3A-C)

500-550 µm. Proximal ridges thin, tall and irregular along the top. Equatorial ridge very smooth and even but with some protrusions along it where the proximal ridges join, both on the proximal and distal side. Ornamentation on both sides is foveolate with weaker walls adjoining the equatorial and proximal ridges. There is no obvious girdle. Surface structure is a very dense cobweb without hairs or loose fibers.

#### *Isoetes capensis* A.V.Duthie ELS96 (Fig. 3D-F)

410-450 µm. Proximal ridges very tall and wide with a small protuberance where they meet. Pulled into very short blunt points where they meet the equatorial ridge. Equatorial ridge slightly lumpy. Ornamentation is pustulate, on the proximal side a few pustules are gathered in the centre of the radial area. On the distal side pustules are irregularly placed and sometimes joining. Girdle with very roughed up fibers and occasionally much smaller pustules. The surface structure is a dense network of fibers with short loose ones.

#### *Isoetes panamensis* Maxon & C.V.Morton ELS112 (Fig. 3G-I)

440-510 µm. Proximal ridges tall, lumpy and pulled into points where they meet the equatorial ridge. Crowning ridge present. Equatorial ridge is very tall and distinctly sinuous. The ornamentation is baculate on both sides with very tall protrusions. No distinguishable girdle. The surface structure is a dense network of fibers with short loose ones.

#### *Isoetes clavata* U.Weber ELS121 (Fig. 3J-L)

410-480 µm. Proximal ridges wide, tall and lumpy with a crowning ridge. Equatorial ridge wide, tall, lumpy and sinuous. The ornamentation is baculate with the protrusions somewhat smaller on the proximal side where they can vary between pustulate and baculate; densely baculate on the distal side. There is no obvious girdle. The surface structure consists of very coarse fibers forming curls, particularly at the summit of the bacules.

#### *Isoetes pedersenii* H.P.Fuchs ex E.I.Meza & Macluf ELS104 (Fig. 4A-C)

410-430 µm. Proximal ridges wide, tall and covered with thick fibers on the sides and with a crowning ridge. Equatorial ridge low, wide and covered with dense fibers on the sides. Ornamentation is on both sides evenly tuberculate with very thick and long fibers radiating from ornamentation and surface. There is a narrow girdle with no tubercles. The surface structure is hairy.

**Figure 4.**
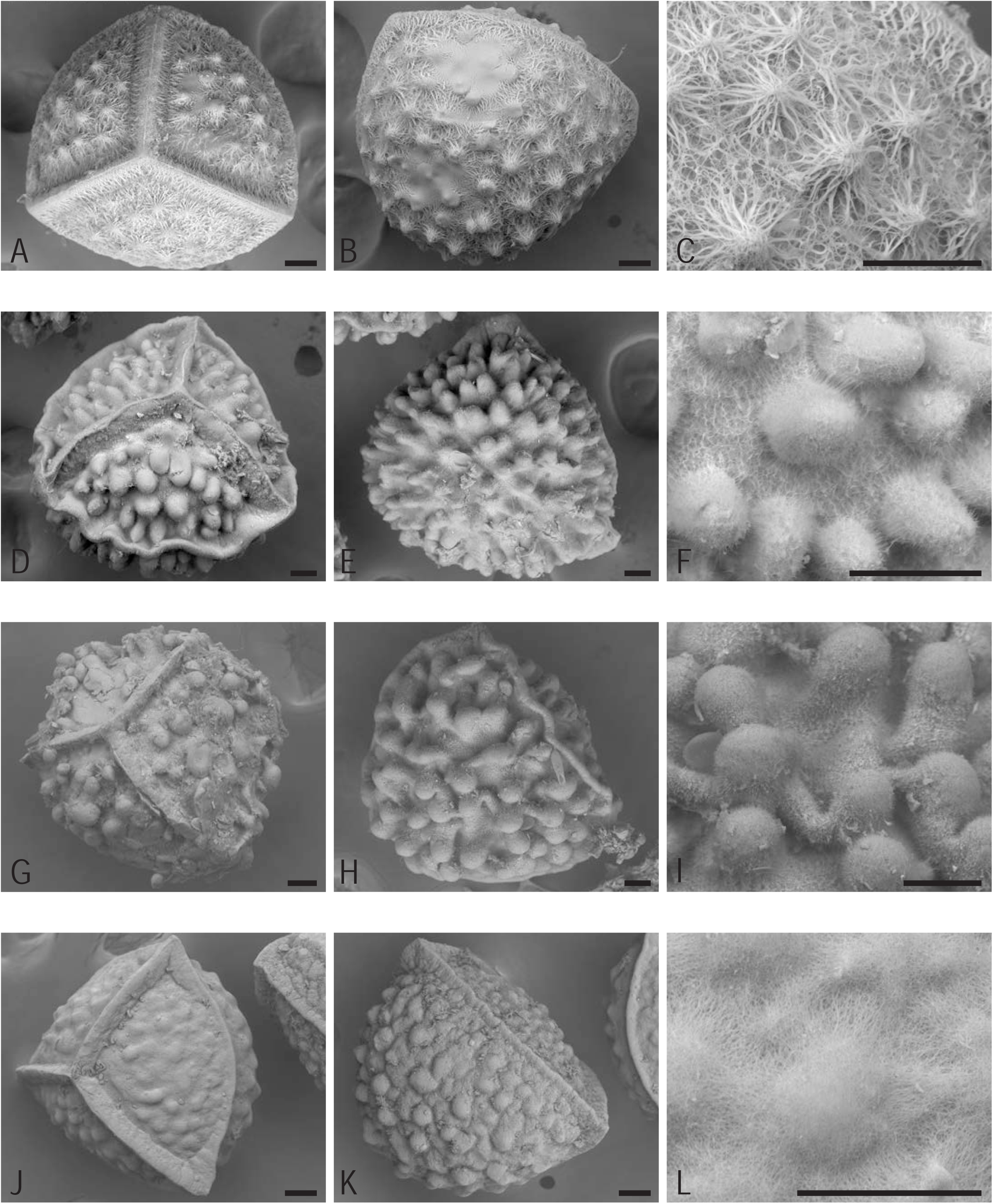
Species of clade A. All scale bars are 50 μm. A-C *Isoetes pedersenii* ELS104, A proximal side, B distal side, C magnified ornamentation on distal side. D-F *Isoetes gardneriana* ELS115, D proximal side, E distal side, F magnified ornamentation on distal side. G-I *Isoetes paraguayensis* ELS111, G proximal side, H distal side, I magnified ornamentation on distal side. J-L *Isoetes melanotheca* ELS03, J proximal side, K distal side, L magnified surface structure on proximal side.

#### *Isoetes gardneriana* Kunze ex A.Braun ELS115 (Fig. 4D-F)

460-520 µm. Proximal ridges very tall, a bit lumpy and pulled into points where they meet the equatorial ridge. Crowning ridge present. Equatorial ridge very tall, sinuous, sometimes lumpy. Ornamentation is on both sides evenly baculate but more densely on the distal side. No obvious girdle. Surface structure is a dense network of fibers with short loose ones.

#### *Isoetes paraguayensis* H.P.Fuchs (nom. nud.) ELS111 (Fig. 4G-I)

400-500 µm. Proximal ridges very tall, thin and lumpy with a crowning ridge. Equatorial ridge tall and sinuous. Ornamentation is evenly baculate on the proximal side, baculate to saccate on the distal side with no obvious girdle. Surface structure is a dense network of fibers with short loose ones.

#### *Isoetes melanotheca* Alston ELS03 (Fig. 4J-L)

400-450 µm. Proximal ridges wide, tall and lumpy with a crowning ridge. Pulled into points where they meet the equatorial ridge. Equatorial ridge thin and sinuous. Ornamentation evenly and densely pustulate apart from an indistinct girdle with folds that gradually turn into pustules. Surface structure is cobweb.

#### *Isoetes coromandelina* subsp. *macrotuberculata* C.R.Marsden ELS72 (Fig. 5A-C)

530-570 µm. Proximal ridges wide, tall, irregular and drawn out into points where they meet the equatorial ridge. Crowning ridge present. Equatorial ridge thin, tall and somewhat sinuous. Ornamentation on proximal side with 1(-4) pustules per radial area, while the distal side has very large tubercles. Surface structure is cobweb.

**Figure 5.**
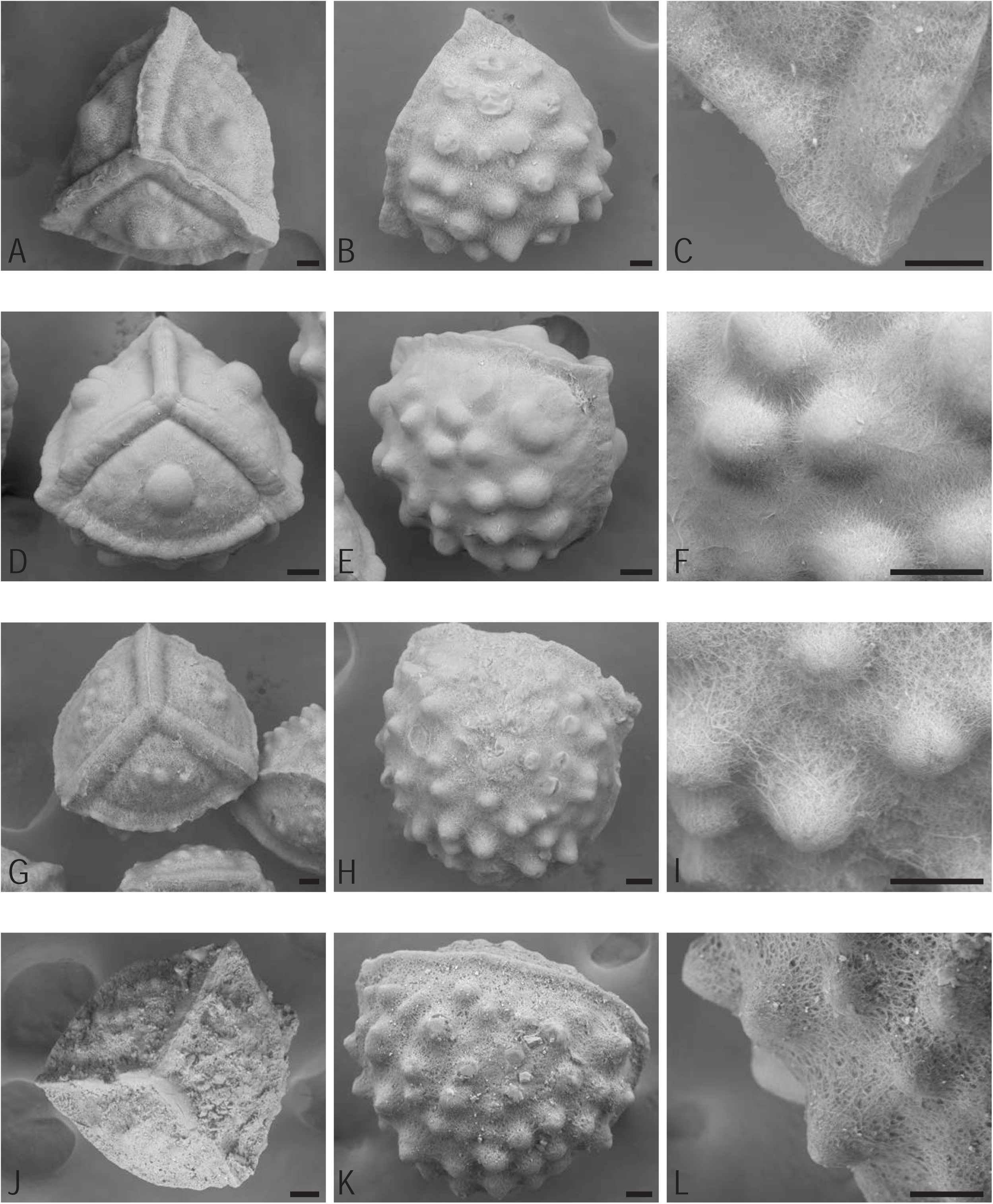
Species of clade A. All scale bars are 50 μm. A-C *Isoetes coromandelina* subsp. *macrotuberculata* ELS72, A proximal side, B distal side, C magnified surface structure on proximal side. D-F *Isoetes coromandelina* ELS117, D proximal side, E distal side, F magnified ornamentation on distal side. G-I *Isoetes pitotii* ELS92, G proximal side, H distal side, I magnified ornamentation on distal side. J-L *Isoetes schweinfurthii* ELS93, J proximal side, K distal side, L magnified ornamentation on distal side.

#### *Isoetes coromandelina* L.f. ELS117 (Fig. 5D-F)

410-440 µm. Proximal ridges wide, tall, a bit lumpy and pulled into points where they meet the equatorial ridge. Crowning ridge present. Equatorial ridge tall and lumpy. Ornamentation pustulate, on proximal side mostly only 1 large pustule per radial area, occasionally flanked by 1-3 smaller ones. On the distal side, evenly distributed large pustules and a girdle with no ornamentation. Surface structure is cobweb with a few course fibers running like veins across the surface.

#### *Isoetes pitotii* Alston ELS92 (Fig. 5G-I)

500-560 µm. Proximal ridges very wide with a crowning ridge. Equatorial ridge narrow, sharp and sinuous. Proximal ornamentation smallish pustules, keeping to the middle of the radial area. On the distal side, even and evenly spread tubercules with a smooth and very narrow girdle. Surface structure is a loose cobweb with constituting fibers congregating near the tubercle points.

#### *Isoetes schweinfurthii* A.Braun ELS93 (Fig. 5J-L)

430-460 µm. Proximal ridges wide with a crowning ridge, pulled into points where they meet the equatorial ridge. Equatorial ridge fairly wide. Ornamentation is tuberculate with a tendency for the protrusions on the proximal side to resemble pustules. A girdle without ornamentation. Surface structure is a cobweb with signs of degradation from age.

#### *Isoetes welwitschii* A.Braun ELS87 (Fig. 6A-C)

470-510 µm. Proximal ridges wide with a crowning ridge. Equatorial ridge thin and sinuous. Ornamentation is tuberculate, on proximal side low tubercles with none at the top of the radial area, on distal side evenly distributed, evenly sized tubercles. Appearing as though they are straining against the surface layer from the inside. A girdle without ornamentation. Surface structure is cobweb.

**Figure 6.**
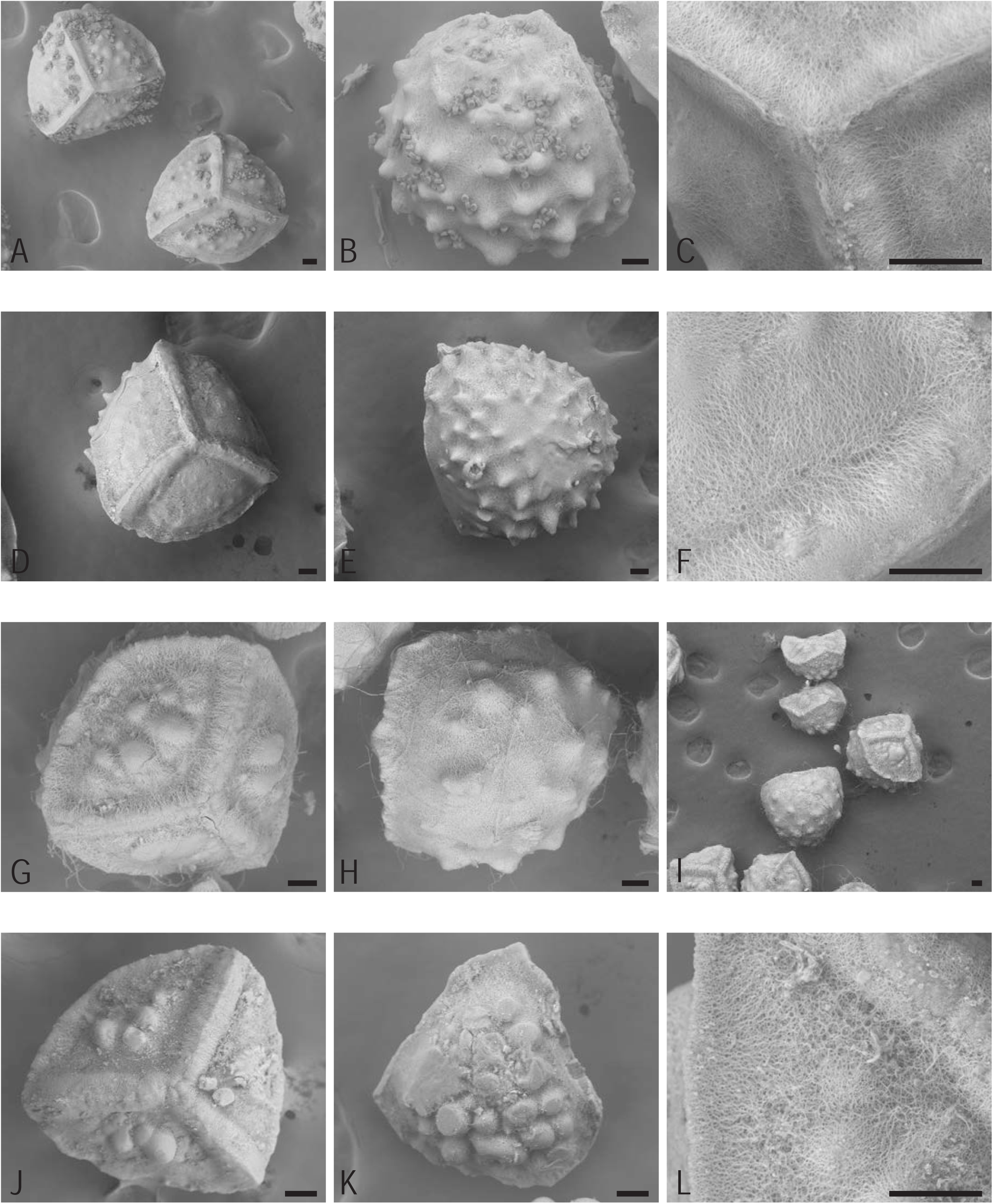
Species of clade A. All scale bars are 50 μm. A-C *Isoetes welwitschii* ELS87, A proximal side, B distal side, C magnified surface structure on proximal side. D-F *Isoetes schweinfurthii* ELS01, D proximal side, E distal side, F magnified surface structure on proximal side. G-I *Isoetes rhodesiana* ELS110, G proximal side, H distal side, I the two spores at the top showing typical appearance of immature spores. J-L *Isoetes schweinfurthii* ELS63, J proximal side, K distal side, L magnified surface structure on proximal side.

#### *Isoetes schweinfurthii* A.Braun ELS01 (Fig. 6D-F)

460-510 µm. Proximal ridges wide with a crowning ridge. Equatorial ridge thin and sinuous. Ornamentation is tuberculate, on proximal side almost completely smooth, the occasional low tubercle, on distal side evenly distributed and evenly sized tubercles. No obvious girdle. Surface structure is cobweb.

#### *Isoetes rhodesiana* Alston ELS110 (Fig. 6G-I)

470-500 µm. Proximal ridges wide and lumpy with a crowning ridge. Equatorial ridge thin and sinuous. Ornamentation is tuberculate. On proximal side few pustules, generally a big one in the middle of the radial area and much smaller ones (if any) on the sides. On the distal side, evenly distributed and evenly sized tubercles. A girdle without ornamentation Surface structure is cobweb.

#### *Isoetes schweinfurthii* A.Braun ELS63 (Fig. 6J-L)

300-450 µm. Proximal ridges wide with a crowning ridge. Equatorial ridge wide and tall. Ornamentation is pustulate. On proximal side few pustules gathered in the middle of the radial area. On distal side evenly distributed pustules. A girdle without ornamentation Surface structure is cobweb to mycelium-like.

#### *Isoetes schweinfurthii* A.Braun ELS02 (Fig. 7A-C)

470-490 µm. Proximal ridges wide with a crowning ridge. Equatorial ridge thin and sinuous. Ornamentation is tuberculate. On proximal side few pustules. On distal side evenly distributed and evenly sized tubercles. A girdle without ornamentation. Surface structure is cobweb.

**Figure 7.**
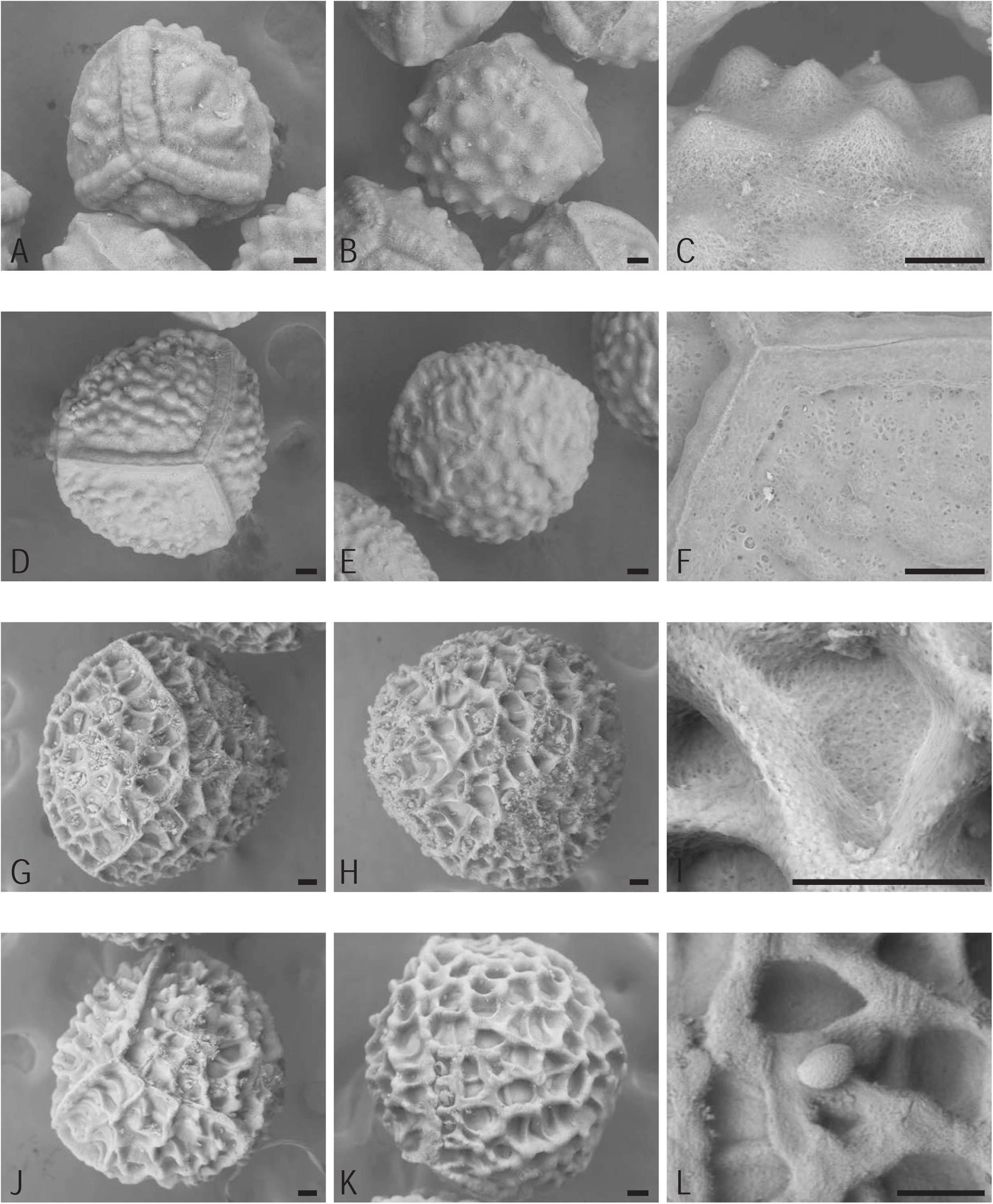
Species of clade A and B. All scale bars are 50 μm. A-C *Isoetes schweinfurthii* ELS02, A proximal side, B distal side, C magnified ornamentation on distal side. D-F *Isoetes histrix* ELS05, D proximal side, E distal side, F magnified surface structure on proximal side. G-I *Isoetes durieui* ELS44, G proximal side, H distal side, I magnified surface structure on distal side. J-L *Isoetes durieui* ELS45, J proximal side, K distal side, L magnified surface structure on distal side with microspore.

#### *Isoetes histrix* Bory ELS05 (Fig. 7D-F)

460-530 µm. Proximal ridges wide with a crowning ridge. Equatorial ridge wavy to lumpy. Ornamentation is pustulate. On proximal side evenly spread pustules, but close to the equatorial ridge there are much smaller ones, more like papillae. On distal side larger pustules, some forming ridges with adjacent ones. No obvious girdle. Surface structure similar to melted cheese with underlying layers visible under an unevenly spread liquid.

#### *Isoetes durieui* Bory ELS44 (Fig. 7G-I)

650-710 µm. Proximal ridges very thin, sharp and tall. Hard to tell apart from the ornamentation. Equatorial ridge thin and tall. Ornamentation is retate. On proximal side retate with taller joints than walls. On distal side retate with taller joints and uneven spaces in between walls. No obvious girdle. Surface structure is sponge-like.

#### *Isoetes durieui* Bory ELS45 (Fig. 7J-L)

640-700 µm. Proximal ridges very thin, sharp and tall. Hard to tell apart from the ornamentation. Equatorial ridge thin and tall. Ornamentation is retate. On proximal side retate with taller joints than walls. On distal side retate with taller joints and uneven spaces in between walls. No obvious girdle. Surface structure is sponge-like.

#### *Isoetes orcuttii* A.A.Eaton ELS51 (Fig. 8A-C)

300-380 µm. Two types of spores in same sporangium, one levigate and smaller the other extremely densely spiculate. Proximal ridges thin, tall and pointed where they meet the equatorial ridge. If spores are spiculate then there are also spicules on the proximal ridges. Equatorial ridge thin and smooth. Ornamentation is spiculate or levigate. On proximal side either very densely covered with spicules or on smaller spores mostly smooth with low pustules near the proximal ridges. On distal side either very densely covered with spicules or on smaller spores levigate. No obvious girdle. Surface structure either overlapping spines or melted cheese.

**Figure 8.**
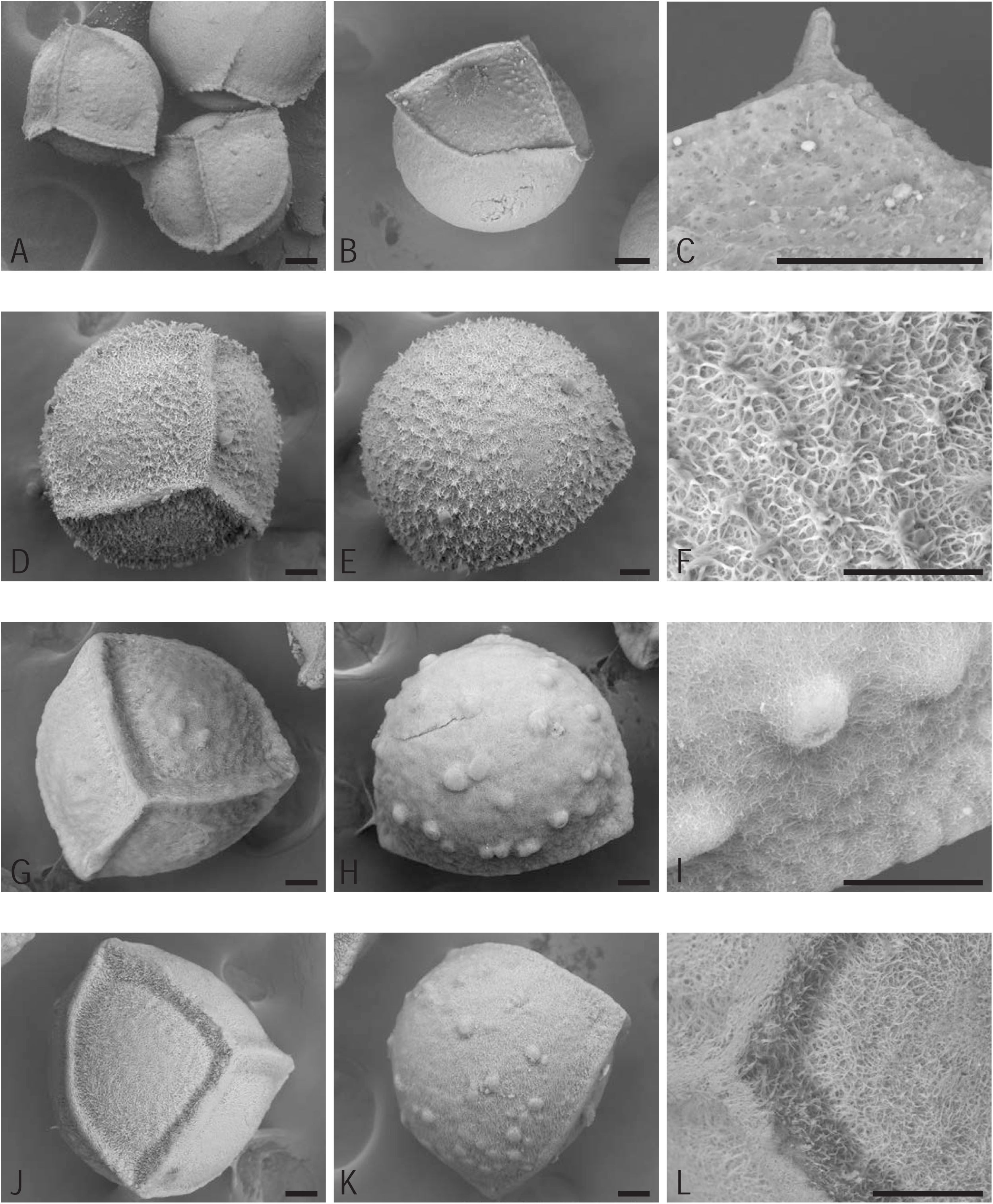
Species of clade B. All scale bars are 50 μm. A-C *Isoetes orcuttii* ELS51, A proximal side, B side, C magnified surface structure on proximal side. D-F *Isoetes nuttallii* ELS53, D proximal side, E distal side, F magnified surface structure on distal side. G-I *Isoetes olympica* ELS88, G proximal side, H distal side, I magnified surface structure on distal side. J-L *Isoetes longissima* subsp. *longissima* ELS17, J proximal side, K distal side, L magnified surface structure on proximal side.

#### *Isoetes nuttallii* A.Braun ex Engelm. ELS53 (Fig. 8D-F)

390-420 µm. Proximal ridges medium wide but tall, crowning ridge not evident on every spore. Equatorial ridge thin and sharp. Ornamentation is spiculate. On both sides spicules made up of wet-looking hairs/fibers, regularly gathering so that the impression from a distance is (inaccurately) of evenly distributed tubercles. No obvious girdle. Surface structure hairy or thorny.

#### *Isoetes olympica* A.Braun ELS88 (Fig. 8G-I)

370-410 µm. Proximal ridges wide, tall and lumpy. Equatorial ridge lumpy. Ornamentation is pustulate. On proximal side very low pustules, towards the middle of the radial area 2-4 slightly larger ones. On distal side unevenly spread pustules. Girdle of smaller pustules. Surface structure network of fibers with short loose ones.

#### Isoetes longissima Bory subsp. longissima ELS17 (Fig. 8J-L)

390-410 µm. Proximal ridges wide and tall. Equatorial ridge wide. Ornamentation is pustulate. On proximal side no obvious ornamentation but the spores are very hairy. On distal side sparsely and unevenly pustulate, distal side tends to be larger than the proximal side. No obvious girdle. Surface structure very hairy.

#### *Isoetes* sp. ELS116 (Fig. 9A-C)

310-390 µm. Proximal ridges wide, tall and pulled into points where they meet the equatorial ridge. Equatorial ridge wide, tall and lumpy to ribbed. Ornamentation is pustulate. On proximal side few pustules gathered in the middle of the radial area. On distal side unevenly spread pustules, exact shape unclear since the spores are so damaged. No obvious girdle. Surface structure very hairy with long, coarse fibers.

**Figure 9.**
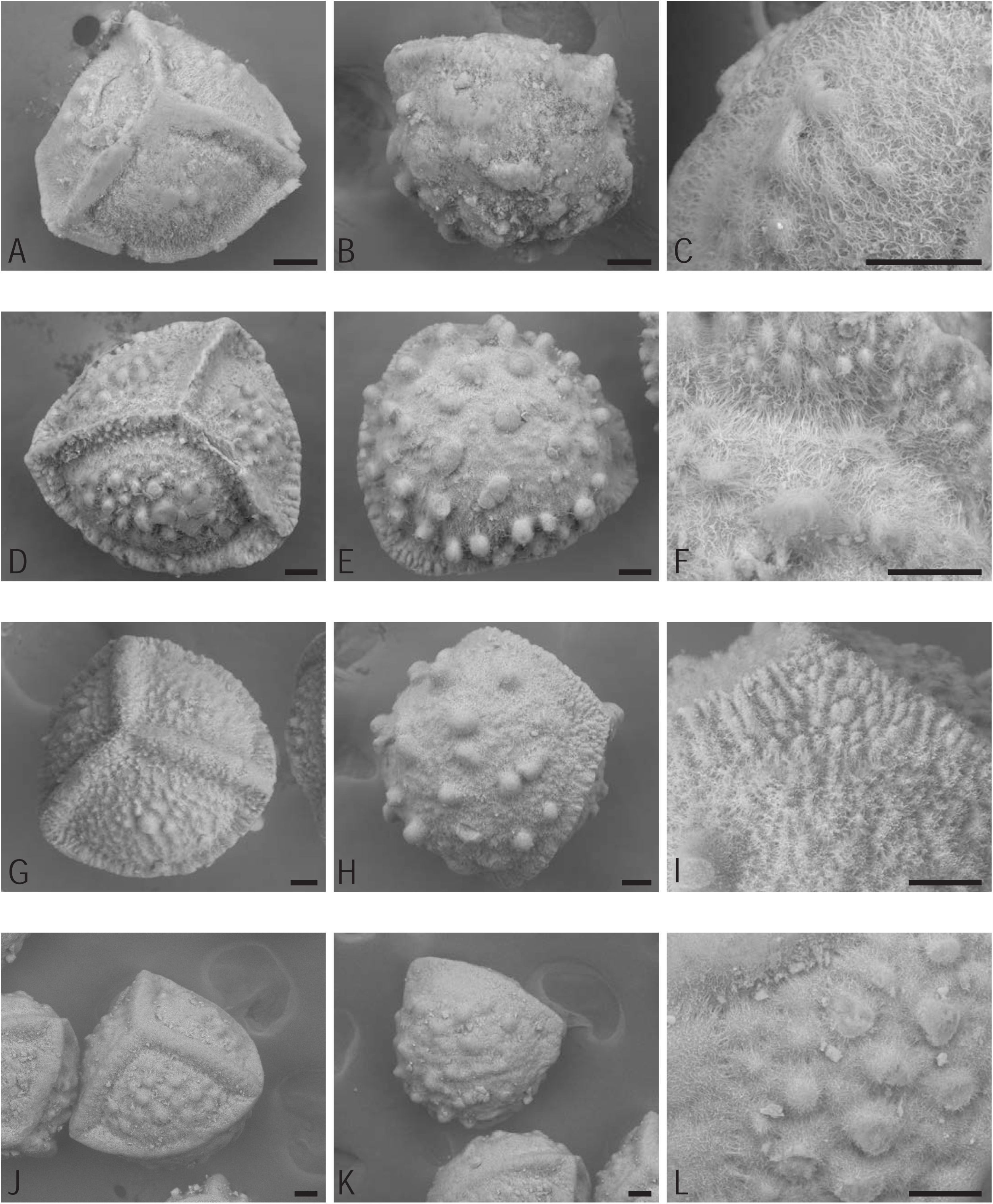
Species of clade B. All scale bars are 50 μm. A-C *Isoetes* sp. ELS116, A proximal side, B distal side, C magnified surface structure on proximal side. D-F *Isoetes dixitii* ELS102, D proximal side, E distal side, F magnified surface structure on proximal side. G-I *Isoetes alstonii* ELS106, G proximal side, H distal side, I magnified surface structure on distal side of the equatorial ridge. J-L *Isoetes giessii* ELS36, J proximal side, K distal side, L magnified ornamentation on proximal side.

#### *Isoetes dixitii* Shende ELS102 (Fig. 9D-F)

400-460 µm. Proximal ridges very tall, a bit lumpy, pulled into points where they meet the equatorial ridge. Ornamentation also on the sides of the ridges. Equatorial ridge very tall and very lumpy, occasionally sinuous. Forming a wide ledge where it meets the proximal ridge. The ledges very ridged on the distal side. Ornamentation is baculate. Radial area covered by smaller tubercles and towards the centre there are bacules, sometimes appearing as though umbellate (but handling damage cannot be ruled out). Everything radiating out long fibers. Distal side varying from densely pustulate to baculate, with the latter being more common. Ornamentation occasionally appearing umbellate (but handling damage cannot be ruled out in these cases). Both ornamentation and surface radiating out long fibers. No obvious girdle. Surface structure coarse, sticky-looking fibers.

#### *Isoetes alstonii* C.F.Reed & Verdc. ELS106 (Fig. 9G-I)

450-500 µm. Proximal ridges wide, tall with a crowning ridge and crowded with ornamentation. Equatorial ridge tall and lumpy to ribbed. Ornamentation is pustulate. Proximal side pustulate and covered in hair/fibers, both from the surface and the ornamentation. Few bigger pustules in the middle of the radial area. Distal side pustulate to baculate as the largest protrusions are quite tall. Protrusions are even and widely spread. There are smaller protrusions underneath the point where the proximal ridges meet the equatorial ridge. Girdle with no pustules. Surface structure is a dense network of fibers with short loose ones.

#### *Isoetes giessii* Launert ELS36 (Fig. 9J-L)

410-450 µm. Proximal ridges wide, tall and with a crowning ridge. Equatorial ridge wide, tall and lumpy to ribbed. Ornamentation is pustulate. Proximal side pustulate and covered in hair/fibers, both from the surface and the ornamentation. On distal side evenly spread pustules. Girdle without pustules. Surface structure hairy.

#### *Isoetes giessii* Launert ELS50 (Fig. 10A-C)

400-470 µm. Proximal ridges wide, tall and with a crowning ridge. Equatorial ridge wide, tall and lumpy to ribbed. Ornamentation is pustulate. Proximal side evenly pustulate, covered in hair/fibers. On distal side evenly spread pustules. No obvious girdle. Surface structure hairy.

**Figure 10.**
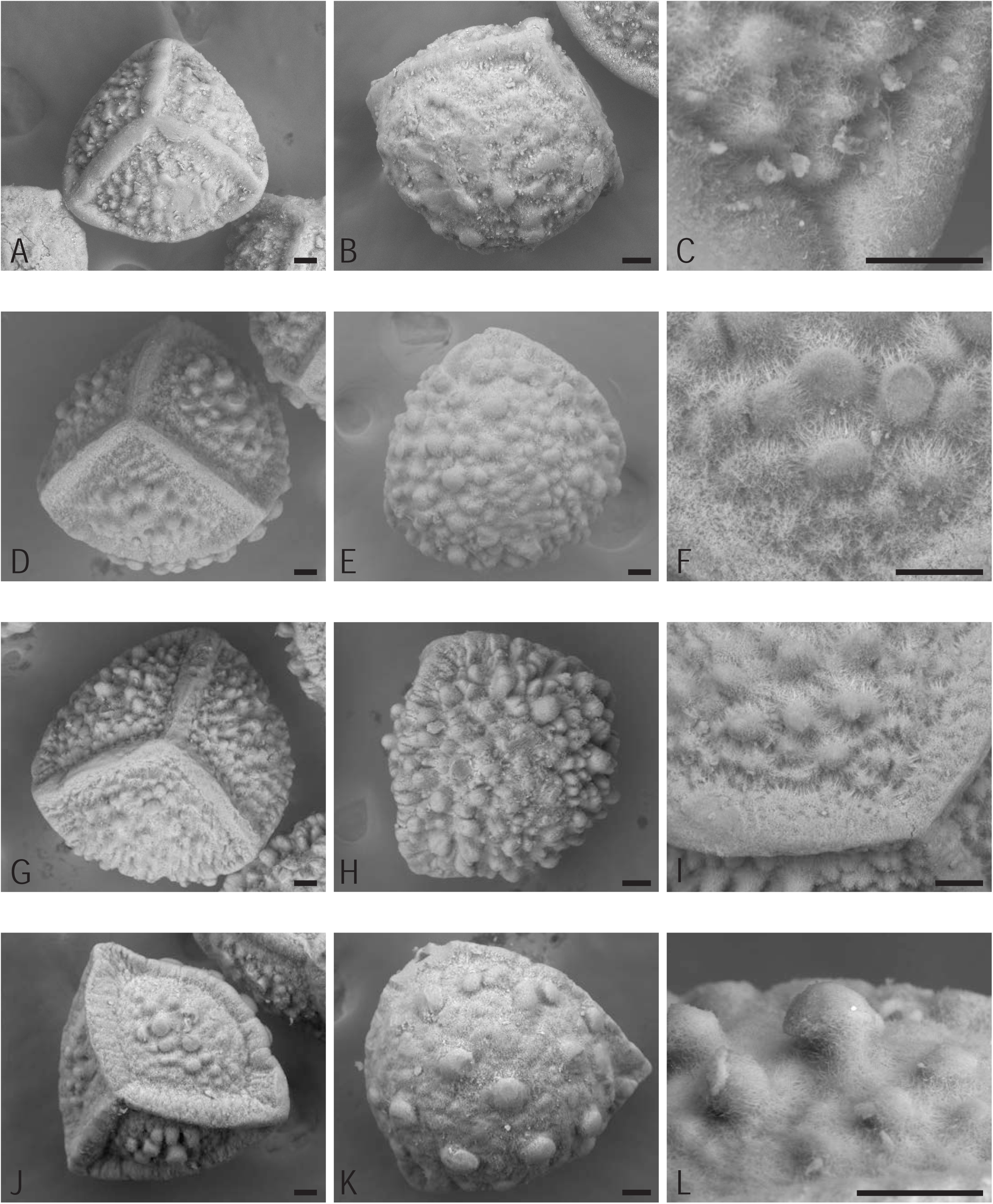
Species of clade B. All scale bars are 50 μm. A-C *Isoetes giessii* ELS50, A proximal side, B distal side, C magnified surface structure on proximal side. D-F *Isoetes* sp. ELS119, D proximal side, E distal side, F magnified surface structure on proximal side. G-I *Isoetes welwitschii* ELS67, G proximal side, H distal side, I magnified surface structure on proximal side. J-L *Isoetes aequinoctialis* ELS54, J proximal side, K distal side, L magnified ornamentation on distal side.

#### *Isoetes* sp. ELS119 (Fig. 10D-F)

530-560 µm. Proximal ridges wide, tall, lumpy with a crowning ridge and pulled into points where they meet the equatorial ridge. Hairy. Equatorial ridge quite wide, tall and lumpy to ribbed. Ornamentation is pustulate. Proximal side evenly pustulate, covered in hairs/fibers. Distal side pustulate to baculate as the largest protrusions are quite tall. Baculate protrusions occasionally cloven. No obvious girdle. Surface structure hairy.

#### *Isoetes welwitschii* A.Braun ELS67 (Fig. 10G-I)

560-590 µm. Proximal ridges wide, tall and with a crowning ridge. Equatorial ridge wide, tall and lumpy to ribbed. Ornamentation is pustulate. Proximal side densely pustulate, covered in hairs/fibers. Distal side pustulate to baculate as the largest protrusions are quite tall. Baculate protrusions occasionally cloven. Girdle without pustules. Surface structure hairy.

#### *Isoetes aequinoctialis* Welw. ex A.Braun ELS54 (Fig. 10J-L)

510-550 µm. Proximal ridges wide, tall, lumpy with a crowning ridge and pulled into points where they meet the equatorial ridge. Equatorial ridge wide, tall and lumpy to ribbed. Ornamentation is umbellate. On proximal side umbellate protrusions in the middle of the radial faces, covered in hairs/fibers. On distal side randomly distributed umbellate protrusions. Covered in hairs/fibers. No obvious girdle. Surface structure hairy.

#### *Isoetes natalensis* Baker ELS113 (Fig. 11A-C)

540-600 µm. Proximal ridges tall, wide and pulled into points where they meet the equatorial ridge. Covered in ornamentation. Equatorial ridge becoming indistinct apart from forming wide ledges where it meets the proximal ridges. Ornamentation is tuberculate. Proximal side densely filled with small tubercles, covered in fibers. On distal side evenly spread small tubercles, covered in fibers. No obvious girdle. Surface structure is a dense network of coarse fibers.

**Figure 11.**
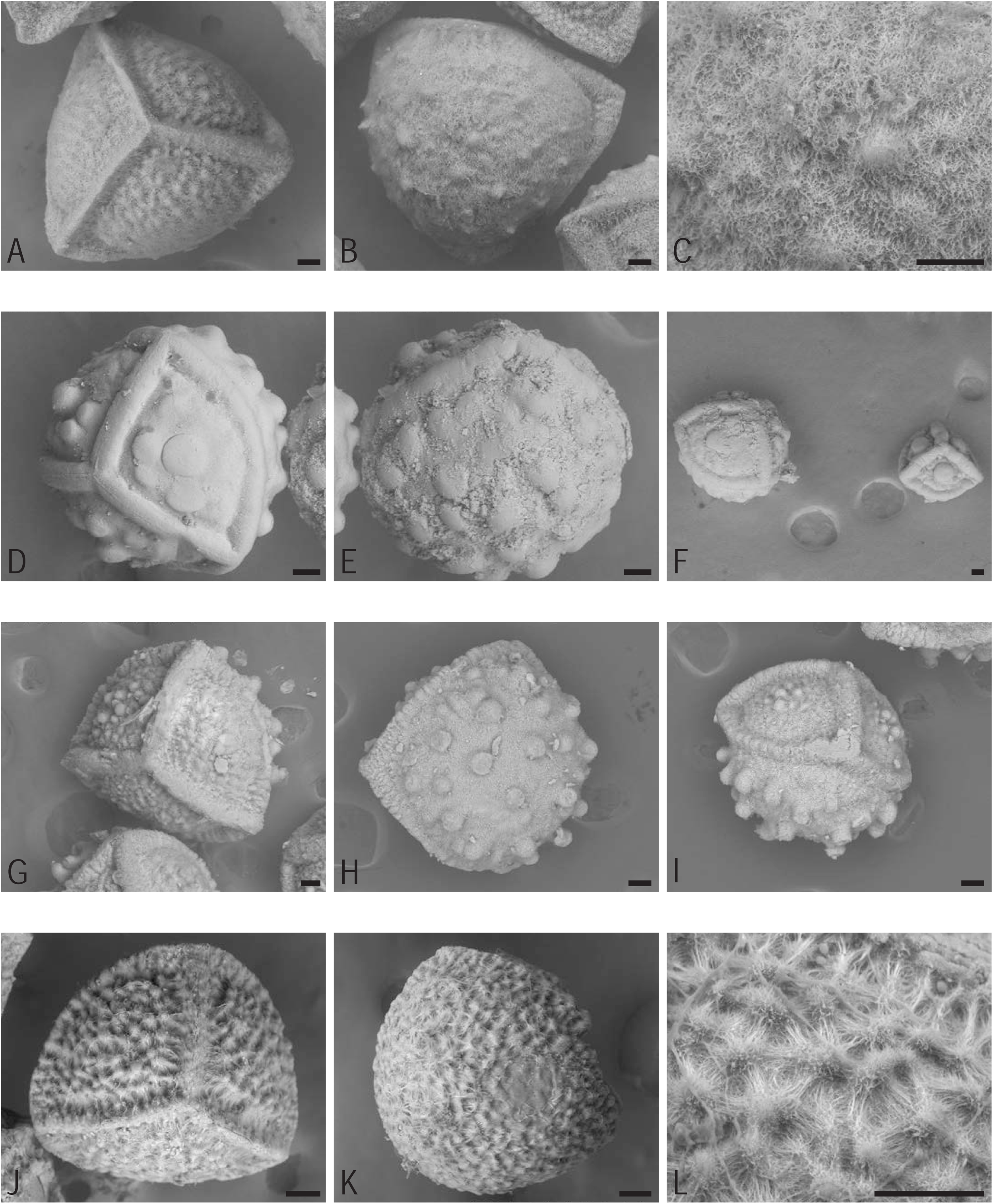
Species of clade B. All scale bars are 50 μm. A-C *Isoetes natalensis* ELS113, A proximal side, B distal side, C magnified surface structure on distal side. D-F *Isoetes aequinoctialis* ELS04, D proximal side, E distal side, F immature spor on the right. G-I *Isoetes aequinoctialis* ELS35, G proximal side, H distal side, I equatorial view showing the girdle. J-L *Isoetes biafrana* ELS46, J proximal side, K distal side, L magnified ornamentation on proximal side.

#### *Isoetes aequinoctialis* Welw. ex A.Braun ELS04 (Fig. 11D-F)

500-570 µm. Proximal ridges wide with a crowning ridge. Equatorial ridge wide. Ornamentation is pustulate. On proximal side few pustules gathered in the middle of the radial area. On distal side evenly distributed pustules. No obvious girdle. Surface structure is unclear.

#### *Isoetes aequinoctialis* Welw. ex A.Braun ELS35 (Fig. 11G-I)

480-590 µm. Proximal ridges wide and tall with a crowning ridge. Equatorial ridge wide, lumpy to ribbed. Ornamentation is pustulate. On proximal side covered in hairs/fibers, both from the surface and the ornamentation. On distal side pustulate to baculate as the largest protrusions are quite tall. Smaller protrusions underneath the point where the proximal ridges meet the equatorial ridge. Girdle without pustules. Surface structure is hairy.

#### *Isoetes biafrana* Alston ELS46 (Fig. 11J-L)

400-420 µm. Proximal ridges wide and tall. Equatorial ridge tall, quite wide and lumpy to sinuous. Ornamentation is pustulate. On proximal side covered in hairs/fibers, both from the surface and the ornamentation. On distal side evenly spread pustules, also on the equatorial ridge. Girdle without pustules. Surface structure is hairy.

#### *Isoetes welwitschii* A.Braun ELS68 (Fig. 12A-C)

430-490 µm. Proximal ridges wide, tall, with indistinct crowning ridge. Ribbed to lumpy and drawn into points where it meets the equatorial ridge. Equatorial ridge wide and tall, lumpy to ribbed, occasionally turning upwards a bit where it meets the proximal ridge so that a pocket forms on either side of each proximal ridge. Ornamentation is umbellate. On proximal side umbellate protrusions in the middle of the radial faces, covered in hairs/coarse fibers. Ornamentation also on the proximal ridge. On distal side randomly distributed umbellate protrusions. Covered in hairs/fibers. Smaller protrusions underneath the point where the proximal ridges meet the equatorial ridge. No obvious girdle. Surface structure is hairy.

**Figure 12.**
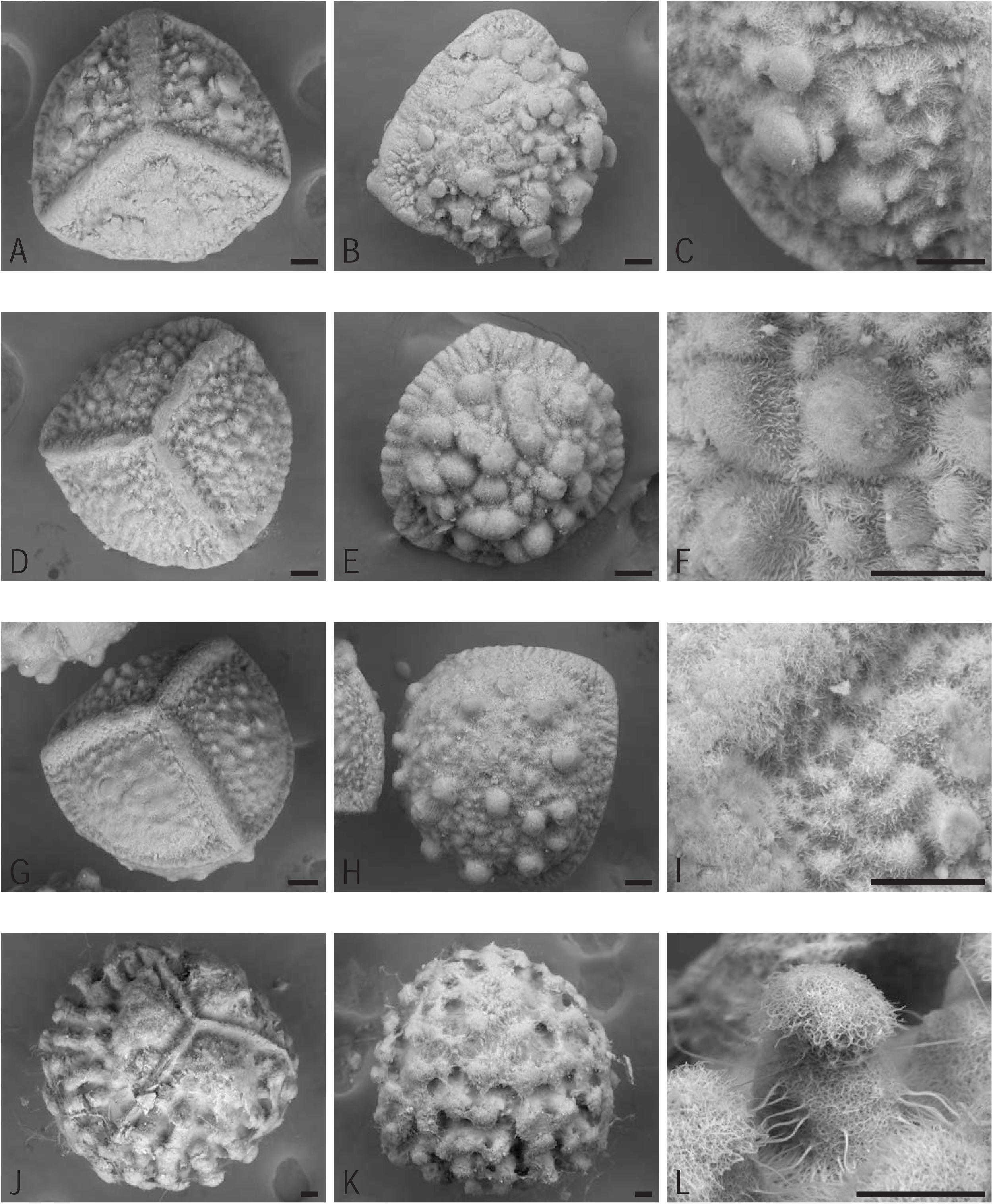
Species of clade B and C. All scale bars are 50 μm. A-C *Isoetes welwitschii* ELS68, A proximal side, B distal side, C magnified ornamentation on proximal side. D-F *Isoetes schweinfurthii* ELS75, D proximal side, E distal side of somewhat immature spore, F magnified surface structure on distal side. G-I *Isoetes welwitschii* ELS33, G proximal side, H distal side, I magnified surface structure on proximal side. J-L *Isoetes malinverniana* ELS126, J proximal side, K distal side, L magnified ornamentation on proximal side.

#### *Isoetes schweinfurthii* A.Braun ELS75 (Fig. 12D-F)

450-480 µm. Proximal ridges wide, tall, uneven, with a crowning ridge. Equatorial ridge lumpy and ribbed to sinuous. Ornamentation is pustulate. On proximal side pustulate and covered in hairs/fibers, both from the surface and the ornamentation. Ornamentation also on the proximal ridge. On distal side pustulate to baculate as the largest protrusions are quite tall. Very hairy. Girdle lacking large protrusions. Surface structure is hairy.

#### *Isoetes welwitschii* A.Braun ELS33 (Fig. 12G-I)

430-460 µm. Proximal ridges wide and tall with a crowning ridge. Equatorial ridge lumpy to ribbed. Ornamentation is pustulate. On proximal side covered in hairs/fibers, both from the surface and the ornamentation. Ornamentation also on the proximal ridge. On distal side pustulate to baculate as the largest protrusions are quite tall. Smaller protrusions underneath the point where the proximal ridges meet the equatorial ridge. Girdle without pustules. Surface structure is hairy.

#### *Isoetes malinverniana* Ces. & De Not. ELS126 (Fig. 12J-L)

700-750 µm (but spores from other viewed specimens are typically smaller, 600-660 µm). Proximal ridges very thin and tall. Equatorial ridge very thin, slightly sinuous and lumpy. Ornamentation is baculate. On proximal side, each radial area has a raised structure closest to the proximal ridge. Below that area are baculate protrusions that are rugged in their upper parts, sometimes adjoined to a raised fibrous netted layer together with other protrusions. On distal side baculate protrusions as on the proximal side but a bit taller and evenly distributed. No obvious girdle. Surface structure is cobweb.

#### *Isoetes japonica* A.Braun ELS114 (Fig. 13A-C)

430-480 µm. Proximal ridges very tall, very narrow and sharp. Equatorial ridge very tall, very narrow, sharp and somewhat upturned. Ornamentation is reticulate. On proximal side reticulate with very tall, narrow and frayed walls. On distal side reticulate with very tall, narrow and frayed walls. No obvious girdle. Surface structure is a network of fibers with pores.

**Figure 13.**
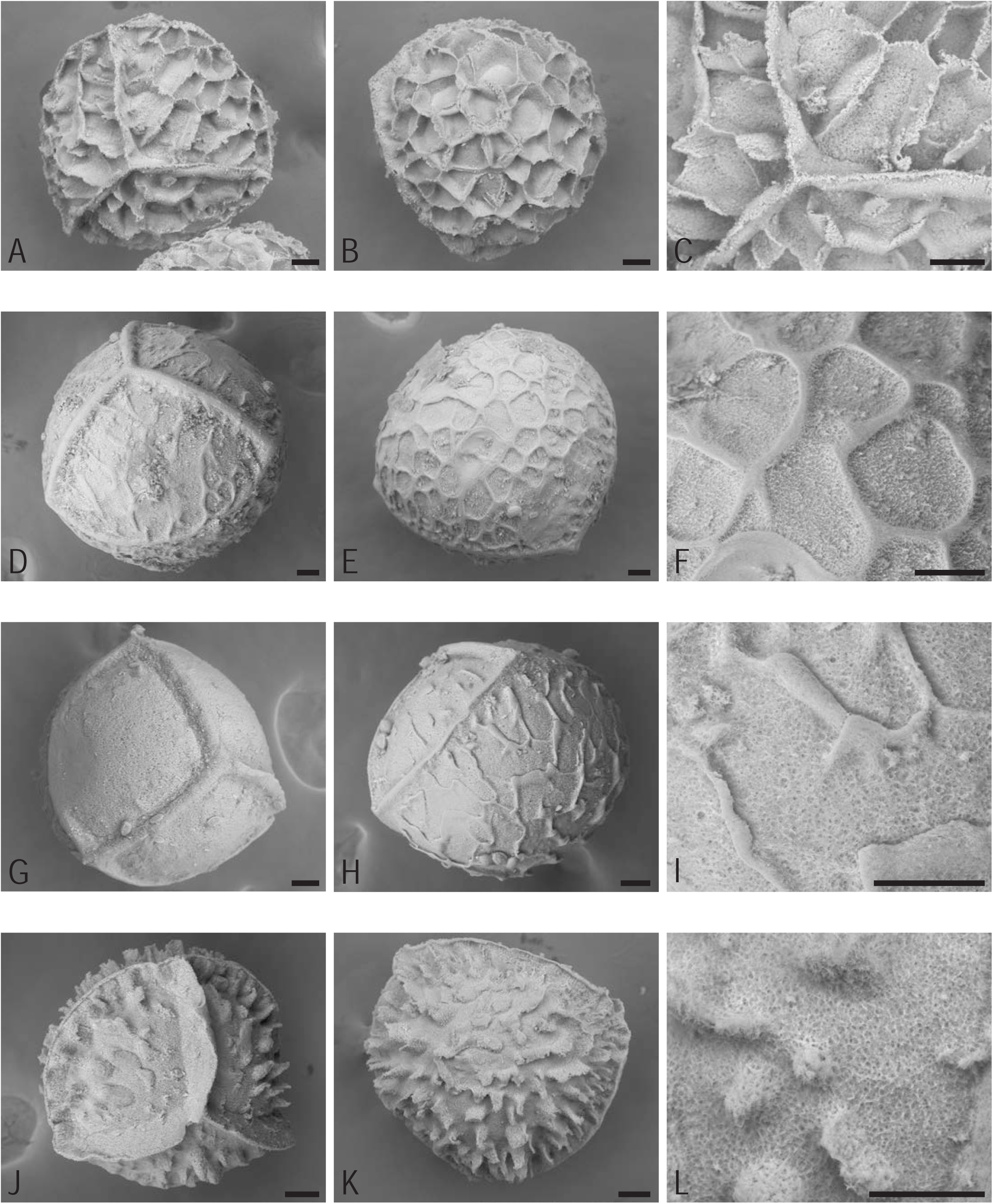
Species of clade D. All scale bars are 50 μm. A-C *Isoetes japonica* ELS114, A proximal side, B distal side, C magnified ornamentation and proximal ridges. D-F *Isoetes stevensii* ELS98, D proximal side, E distal side with microspore, F magnified surface structure on distal side. G-I *Isoetes philippinensis* ELS94, G proximal side, H distal side with microspores, I magnified surface structure on distal side. J-L *Isoetes sinensis* ELS109, J proximal side, K distal side, L magnified surface structure on proximal side.

#### *Isoetes stevensii* J.R.Croft ELS98 (Fig. 13D-F)

570-640 µm. Proximal ridges quite low and smooth on top. Equatorial ridge very low, roughly the same height as the ornamentation. Ornamentation is retate. On both sides retate with low, wide walls, appearing as though smoothed down. They often butt out over the areoles, forming concave walls. Areoles covered by sticky-looking fibers. No obvious girdle. Surface structure is a dense network of fibers with short loose ones.

#### *Isoetes philippinensis* Merr. & L.M.Perry ELS94 (Fig. 13G-I)

450-480 µm. Proximal ridges tall and pulled into points where they meet the equatorial ridge. Equatorial ridge narrow and turned upwards. Ornamentation is rugulate. On proximal side either smooth or with short ridges all arranged closer to the equatorial ridge and perpendicular to it. On distal side rugulate with walls of irregular height folded over, as if in strong wind. Rugulate or reticulate, but the walls are not closing as often as in reticulate spores of other species. No obvious girdle. Surface structure is a dense network of fibers with short loose ones.

#### *Isoetes sinensis* T.C.Palmer ELS109 (Fig. 13J-L)

370-410 µm. Proximal ridges very tall, very narrow and sharp. Equatorial ridge very tall, very narrow, sharp and somewhat directed upwards. Ornamentation is echinate. On distal side echinate to almost reticulate on some spores. No obvious girdle. Surface structure is a network of fibers with pores.

#### *Isoetes drummondii* A.Braun ELS27 (Fig. 14A-C)

410-480 µm. Proximal ridges thin with a crowning ridge. Equatorial ridge narrow, sharp and somewhat uneven. Ornamentation is tuberculate. On proximal side covered in hairs/fibers, both from the surface and the ornamentation. Ornamentation also on the proximal ridge. On distal side tuberculate and covered in hairs/fibers. Some tubercles occasionally joining to form short ridges. Girdle with much smaller tubercles. Surface structure is hairy.

**Figure 14.**
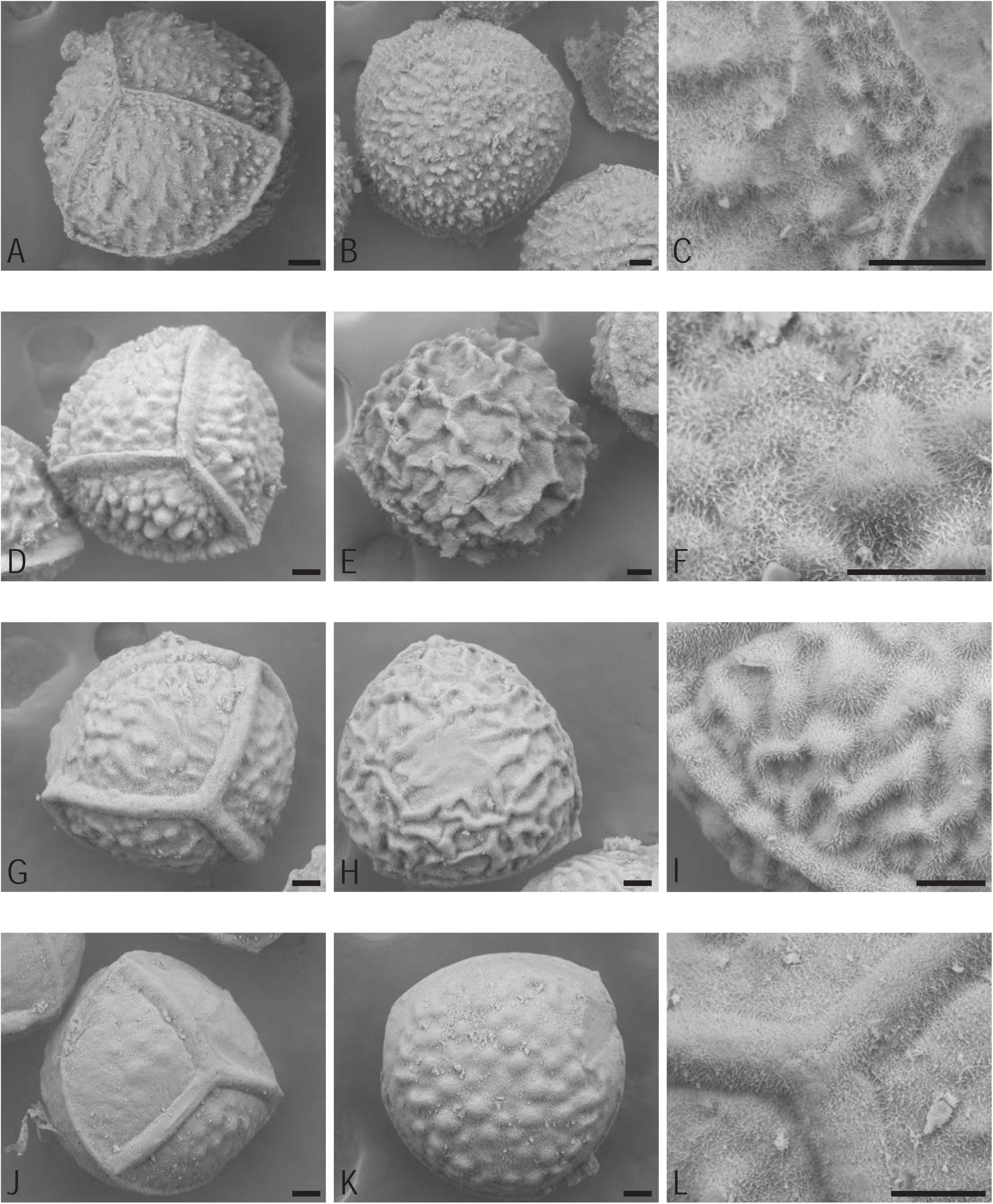
Species of clade D. All scale bars are 50 μm. A-C *Isoetes drummondii* ELS27, A proximal side, B distal side, C magnified ornamentation and proximal ridges. D-F *Isoetes sampathkumaranii* ELS49, D proximal side, E distal side, F magnified surface structure on proximal side. G-I *Isoetes muelleri* ELS105, G proximal side, H distal side, I magnified ornamentation on proximal side and equatorial ridge. J-L *Isoetes kirkii* ELS78, J proximal side, K distal side, L magnified surface structure and proximal ridges.

#### *Isoetes sampathkumaranii* L.N.Rao ELS49 (Fig. 14D-F)

410-490 µm. Proximal ridges wide, tall, with a crowning ridge and forming points when meeting the equatorial ridge. Equatorial ridge very tall, sinuous. Ornamentation is pustulate to rugulate. On both sides rugulate with tall folds, covered in hairs/fibers. No obvious girdle. Surface structure is hairy.

#### *Isoetes muelleri* A.Braun ELS105 (Fig. 14G-I)

460-510 µm. Proximal ridges wide, pulled into points where they meet the equatorial ridge. Very hairy. Equatorial ridge tallish, lumpy, a bit of overhang on the distal side. Ornamentation is rugulate. On proximal side rugulate tending to be ridges. Very hairy. On distal side rugulate in wide folds. Very hairy. No obvious girdle. Surface structure is hairy.

#### *Isoetes kirkii* A.Braun ELS78 (Fig. 14J-L)

470-530 µm. Proximal ridges wide, no ornamentation on them. Equatorial ridge thin and low. Ornamentation is pustulate. On proximal side very low pustules that are covered in short fibers. Larger pustules towards the top of the radial areas. On distal side low pustules that are covered in short fibers. Girdle without ornamentation. Surface structure is hairy.

#### *Isoetes humilior* F.Muell. ex A.Braun ELS69 (Fig. 15A-C)

550-630 µm. Proximal ridges wide, with a crowning ridge and with a low nib where they join in the middle. Equatorial ridge smooth, with a thin overhang towards the distal side. Ornamentation is rugulate. On proximal side rugulate to pustulate with the top of the ornamentation flattened out. On distal side mostly pustulate, but some have joined up becoming rugulate, again the ornamentation is flattened on the top. Girdle without ornamentation. Surface structure fibers on the surface, below risen dough.

**Figure 15.**
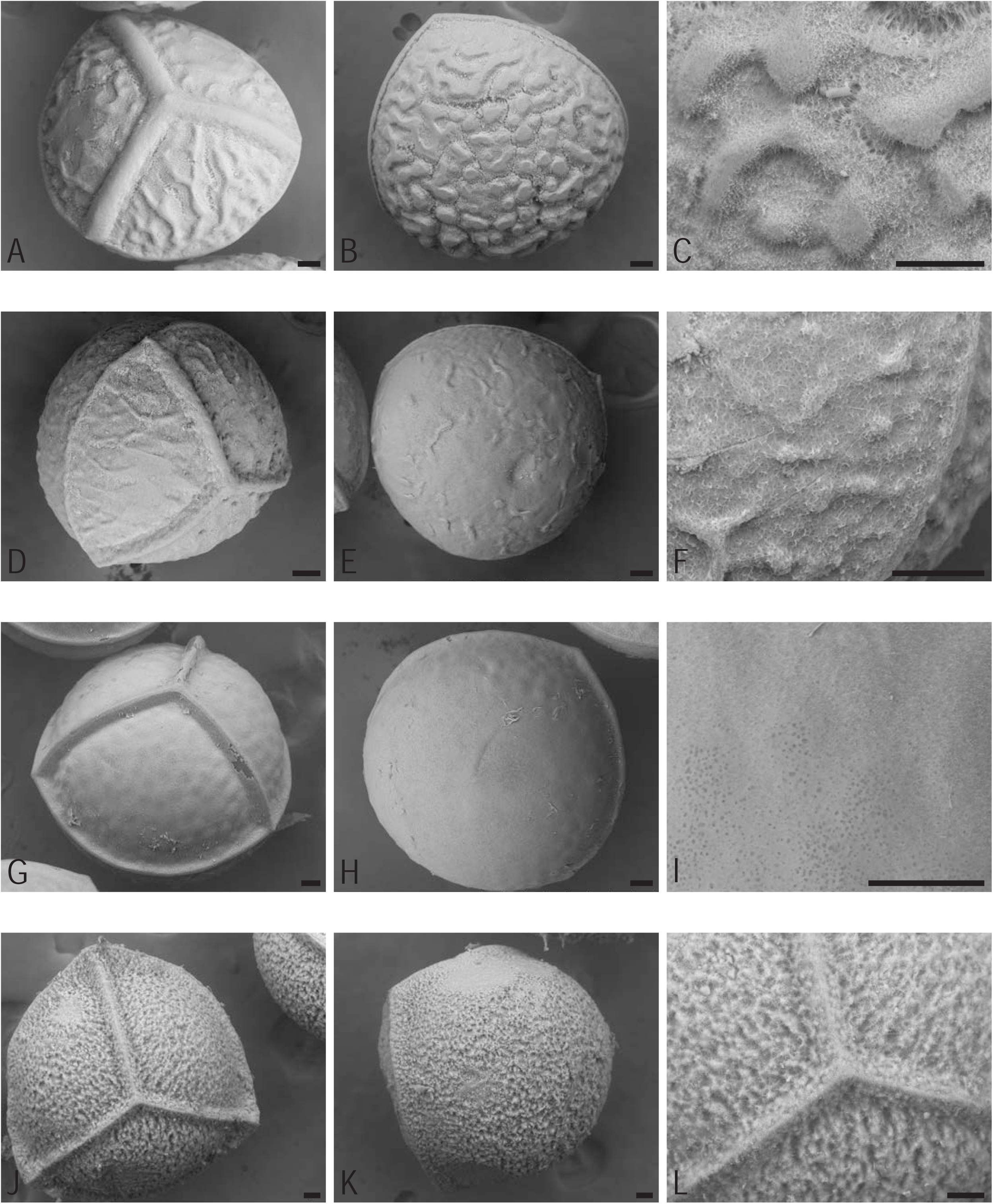
Species of clade D and E. All scale bars are 50 μm. A-C *Isoetes humilior* ELS69, A proximal side, B distal side, C magnified ornamentation on distal side. D-F *Isoetes alpina* ELS100, D proximal side, E distal side, F magnified ornamentation on proximal side and equatorial ridge. G-I *Isoetes andicola* ELS108, G proximal side, H distal side, I magnified surface structure on proximal side. J-L *Isoetes andina* ELS103, J proximal side, K distal side, L magnified surface structure and proximal ridges.

#### *Isoetes alpina* Kirk ELS100 (Fig. 15D-F)

490-510 µm. Proximal ridges wide, lumpy with a crowning ridge. Tall upper ridge close to the points. Equatorial ridge low and narrow with an overhang towards the distal side. Ornamentation is rugulate. On proximal side indistinctly rugulate with a few unevenly spaced tubercules towards the equatorial ridge. On distal side very weakly rugulate with few ridges standing a little taller. Girdle with low undulations/pustules. Surface structure hairy, a network of fibers with short loose ones and pores.

#### *Isoetes andicola* (Amstutz) L.D.Gómez ELS108 (Fig. 15G-I)

580-610 µm. Proximal ridges tall, medium wide and very smooth. Equatorial ridge low and very smooth. Ornamentation is levigate. On proximal side slightly undulating surface with short hair-like fibers near the proximal ridges. On distal side slightly undulating surface with short hair-like fibers near the equatorial ridge. No obvious girdle. Surface structure dense with pores.

#### *Isoetes andina* Kirk ELS103 (Fig. 15J-L)

760-780 µm. Proximal ridges narrow, low with a crowning ridge. Pulled into points where they meet the equatorial ridge. Ornamentation continues over it. Equatorial ridge tall and a little bit wider than the proximal ridges. Ornamentation continues over it. Ornamentation is cristate. On proximal side evenly cristate. Hairy. On distal side evenly cristate. Hairy. Larger distal side than proximal. No obvious girdle. Surface structure is hairy.

#### *Isoetes mexicana* Underw. ELS84 (Fig. 16A-C)

410-460 µm. Proximal ridges wide with a crowning ridge and no ornamentation on them. Equatorial ridge very tall, thin, slightly sinuous. Ornamentation is tuberculate. On proximal side evenly tuberculate. Extremely hairy, both from the ornamentation and the surface. On distal side densely tuberculate, very hairy, both from the ornamentation and the surface. Girdle without ornamentation or the odd low tubercule. Surface structure is hairy.

**Figure 16.**
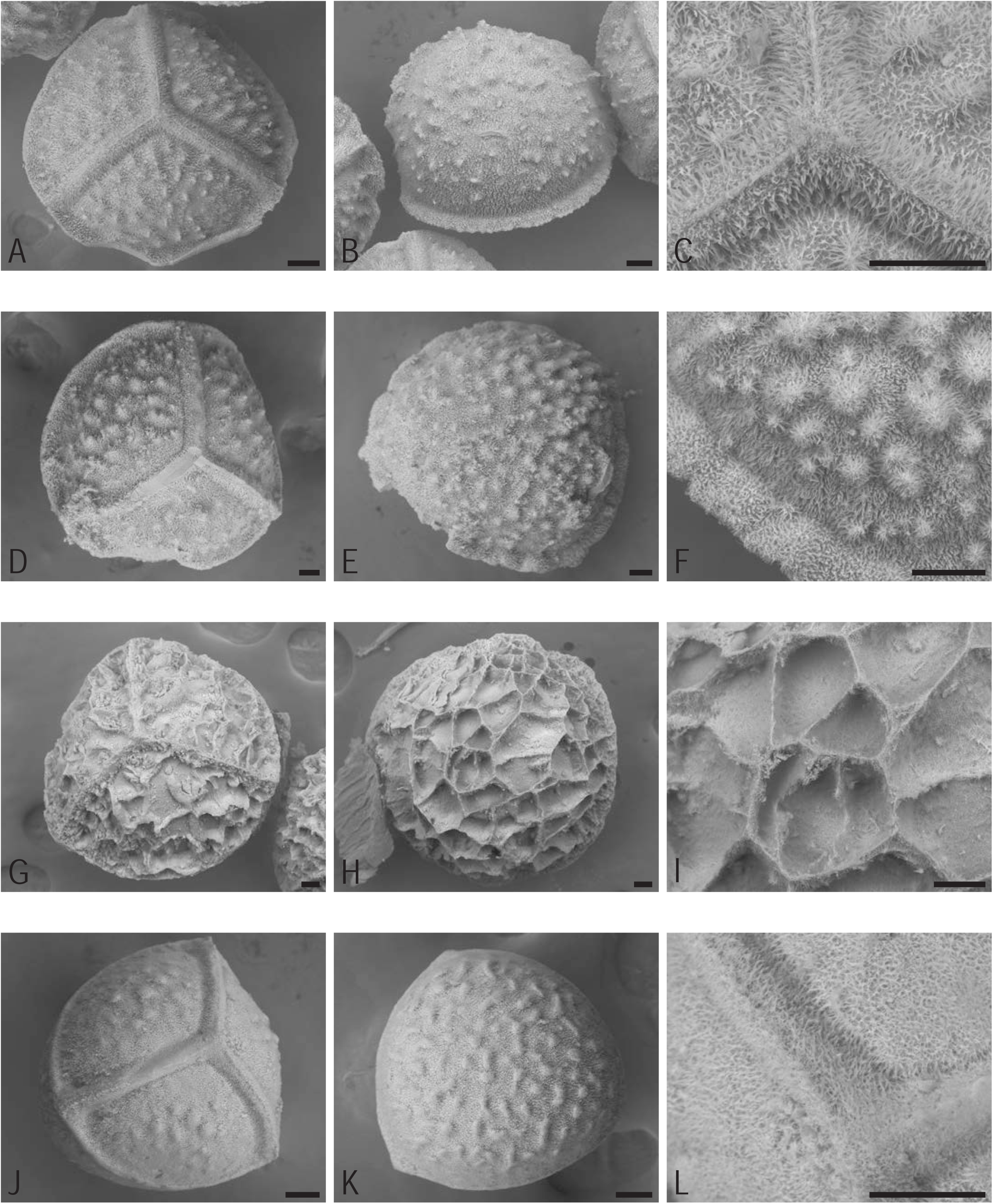
Species of clade E. All scale bars are 50 μm. A-C *Isoetes mexicana* ELS84, A proximal side, B distal side, C magnified surface structure and proximal ridges. D-F *Isoetes montezumae* ELS83, D proximal side, E distal side, F magnified ornamentation on distal side. G-I *Isoetes novo-granadensis* ELS31, G proximal side, H distal side, I magnified ornamentation on distal side. J-L *Isoetes bolanderi* ELS118, J proximal side, K distal side, L magnified surface structure and proximal ridges.

#### *Isoetes montezumae* A.A.Eaton ELS83 (Fig. 16D-F)

410-500 µm. Proximal ridges wide with a crowning ridge and no ornamentation on them. Equatorial ridge very tall, thin and slightly sinuous. Ornamentation is tuberculate. On proximal side evenly tuberculate. Extremely hairy, both from the ornamentation and the surface. On distal side densely tuberculate, very hairy, both from the ornamentation and the surface. Girdle without ornamentation or the odd low tubercule. Surface structure is hairy.

#### *Isoetes novo-granadensis* H.P.Fuchs ELS31 (Fig. 16G-I)

610-710 µm. Proximal ridges very thin and tall with cristate protrusions along the sides. Equatorial ridge thin and sharp. Ornamentation is reticulate. On proximal side tall thin walls. Small protrusions along the proximal ridges with fibers emanating from them. On distal side thin tall walls with fibers emanating from the top of them. Unevenly sized spaces. No obvious girdle. Surface structure is dense with a network of thin fibers as outmost layer.

#### *Isoetes bolanderi* Engelm. ELS118 (Fig. 16J-L)

350-440 µm. Proximal ridges wide with a crowning ridge and pulled into points where they meet the equatorial ridge. Hairy. Equatorial ridge low and hairy. Ornamentation is pustulate. On proximal side low pustules, all covered by dense fur. On distal side evenly spread pustules, sometimes joined. Covered in hairs. Distal half often seemingly flattened along the bottom, not forming an even half-orb. Wide girdle without ornamentation. Surface structure is hairy.

#### *Isoetes flaccida* var. *chapmanii* Engelm. ELS107 (Fig. 17A-C)

440-540 µm. Proximal ridges wide-ish, low and pulled into points where they meet the equatorial ridge. Equatorial ridge wide and smooth. Ornamentation is rugulate. Proximal side smooth or hints of very low pustules. On distal side very low rugulate folds or pustules that appear shaved off along the top. Dense network of fibers between them. No obvious girdle. Surface structure is a network of fibers with pores.

**Figure 17.**
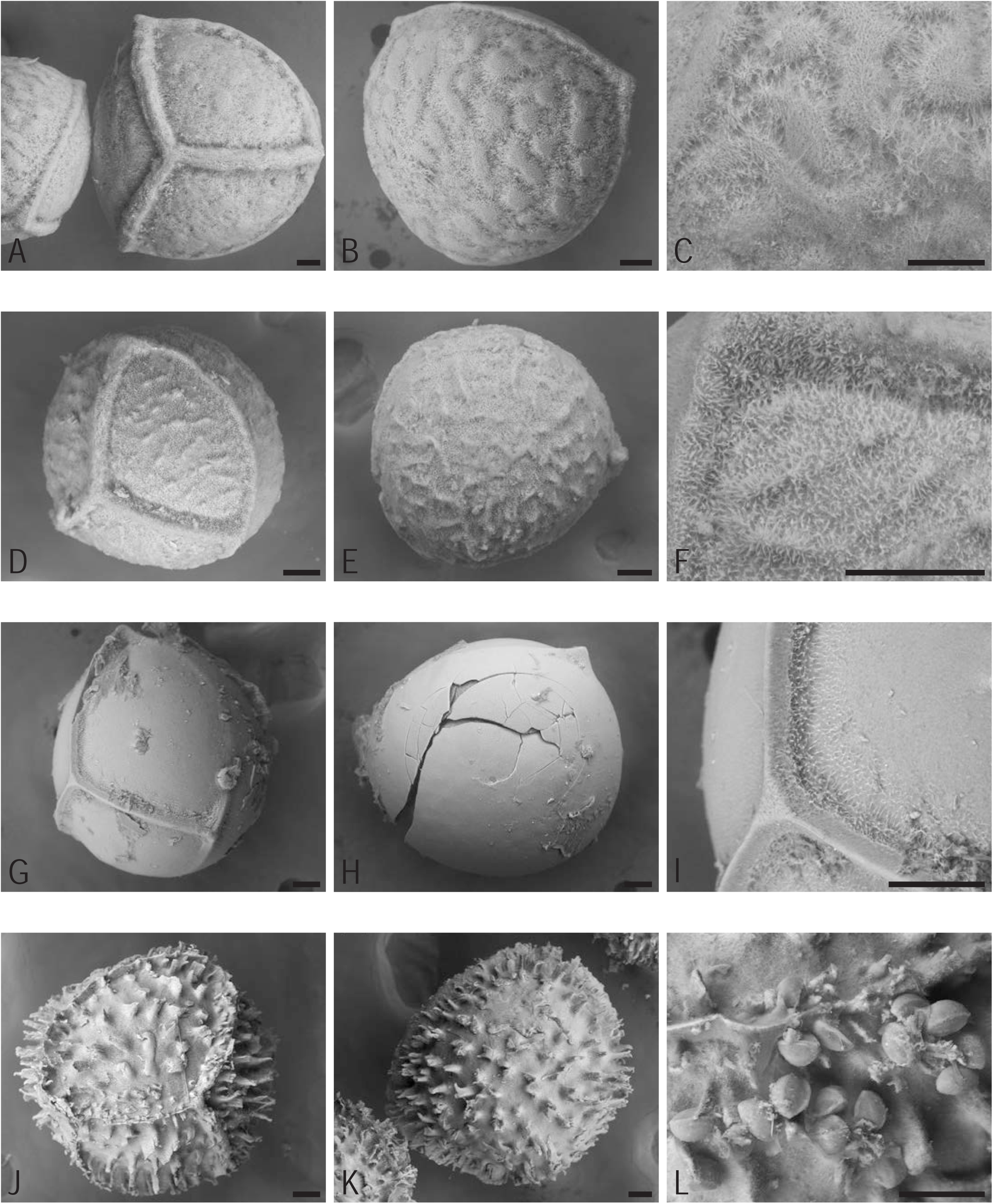
Species of clade E. All scale bars are 50 μm. A-C *Isoetes flaccida* var. *chapmanii* ELS107, A proximal side, B distal side, C magnified ornamentation on distal side. D-F *Isoetes melanopoda* ELS99, D proximal side, E distal side, F magnified surface structure on proximal side, proximal ridge above and equatorial ridge to the left. G-I *Isoetes palmeri* ELS32, G proximal side, H distal side, I magnified surface structure on proximal side. J-L *Isoetes riparia* ELS85, J proximal side, K distal side, L microspores just below the equatorial ridge.

#### *Isoetes melanopoda* J.Gay & Durieu ELS99 (Fig. 17D-F)

360-380 µm. Proximal ridges wide-ish, of medium height, not smooth. Equatorial ridge fairly narrow, tendency to overhang on the distal side. Ornamentation is rugulate. On proximal side in usual rugulate style there are short ridges perpendicular to the equatorial ridge. On distal side rugulate but hard to see the exact form of the ridge because it is very hairy. Girdle without ornamentation. Surface structure is hairy.

#### *Isoetes palmeri* H.P.Fuchs ELS32 (Fig. 17G-I)

480-510 µm. Proximal ridges thin-ish and pointed where they meet the equatorial ridge. Equatorial ridge thin, low, almost disappearing along the radial faces. Ornamentation is levigate. On proximal side levigate but with some fibers alongside the proximal ridges. On distal side levigate but with some fibers under the points where the proximal ridges meet the equatorial ridge. No obvious girdle. Surface structure is dense.

#### *Isoetes riparia* Engelm. ex A.Braun ELS85 (Fig. 17J-L)

460-540 µm. Proximal ridges very thin, tall. Ornamentation also on the ridges. Equatorial ridge very thin, sharp and tall. Ornamentation is echinate. On both sides protrusions are sometimes joining into broader bands rather than solitary needles. No obvious girdle. Surface structure is dense with a network of thin fibers as outmost layer.

#### *Isoetes* sp. ELS52 (Fig. 18A-C)

460-490 µm. Proximal ridges tall and medium wide with a crowning ridge, appearing quite smooth but occasionally carrying ornamentation. Equatorial ridge very thin, sharp and tall. Ornamentation is echinate. On both sides protrusions are sometimes joining into broader bands rather than solitary needles. No obvious girdle. Surface structure is dense and slightly grainy.

**Figure 18.**
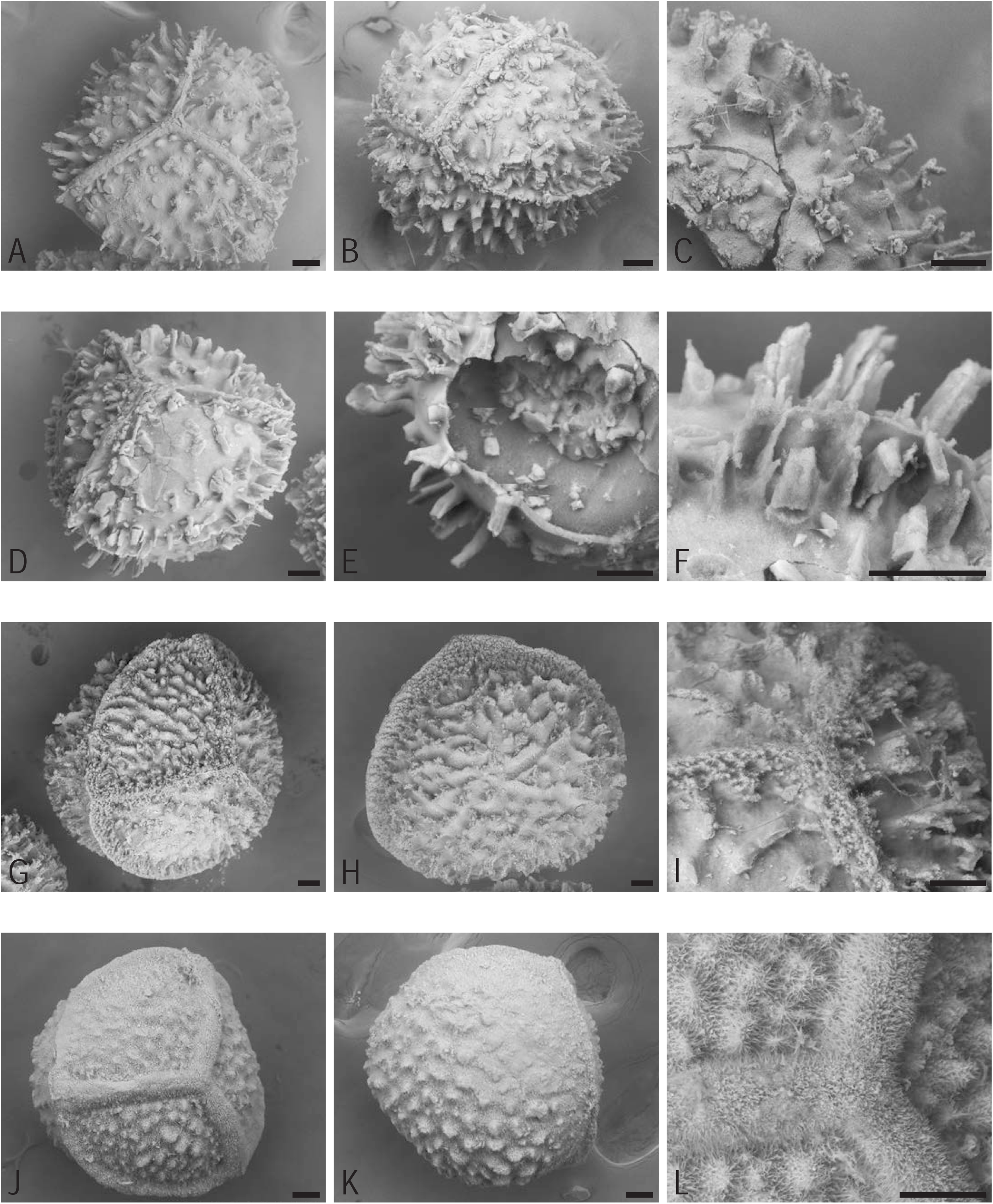
Species of clade E. All scale bars are 50 μm. A-C *Isoetes* sp. ELS52, A proximal side, B side, C magnified ornamentation on distal side. D-F *Isoetes* sp. ELS55, D proximal side, E distal side of a collapsed spore, F magnified ornamentation on proximal side and equatorial ridge. G-I *Isoetes saccharata* ELS73, G proximal side, H distal side, I magnified ornamentation and proximal ridges. J-L *Isoetes howellii* ELS77, J proximal side, K distal side, L magnified ornamentation and proximal ridges.

#### *Isoetes* sp. ELS55 (Fig. 18D-F)

390-420 µm. Proximal ridges tall and medium wide with a crowning ridge, occasionally carrying ornamentation. Equatorial ridge very thin, sharp and tall. Ornamentation is echinate. On both sides protrusions are sometimes joining into broader bands rather than solitary needles. No obvious girdle. Surface structure is dense with a network of thin fibers as outmost layer.

#### *Isoetes saccharata* Engelm. ELS73 (Fig. 18G-I)

580-660 µm. Proximal ridges very thin, tall and completely encrusted by small protrusions. Equatorial ridge very thin, sharp and tall. Ornamentation is echinate. On proximal side the protrusions are sometimes joining into broader bands rather than solitary needles. Shorter spines as compared to those of *I. echinospora*. On distal side echinate to rugulate where the spines join to form short walls. Girdle of densely cristate protrusions. Surface structure is dense with a network of thin fibers as outmost layer.

#### *Isoetes howellii* Engelm. ELS77 (Fig. 18J-L)

470-510 µm. Proximal ridges wide with a crowning ridge, no ornamentation on them. Equatorial ridge regular, can appear thin from the distal side. Ornamentation is tuberculate. On proximal side evenly tuberculate and very hairy, both from the ornamentation and the surface. On distal side densely tuberculate and very hairy, both from the ornamentation and the surface. Girdle without ornamentation. Surface structure is hairy.

#### *Isoetes lechleri* Mett. ELS42 (Fig. 19A-C)

400-450 µm. Proximal ridges wide, tall and forming swollen points when meeting the equatorial ridge. Equatorial ridge of medium height and even. Ornamentation is levigate. On both sides devoid of features. No obvious girdle. Surface structure is dense with pores.

**Figure 19.**
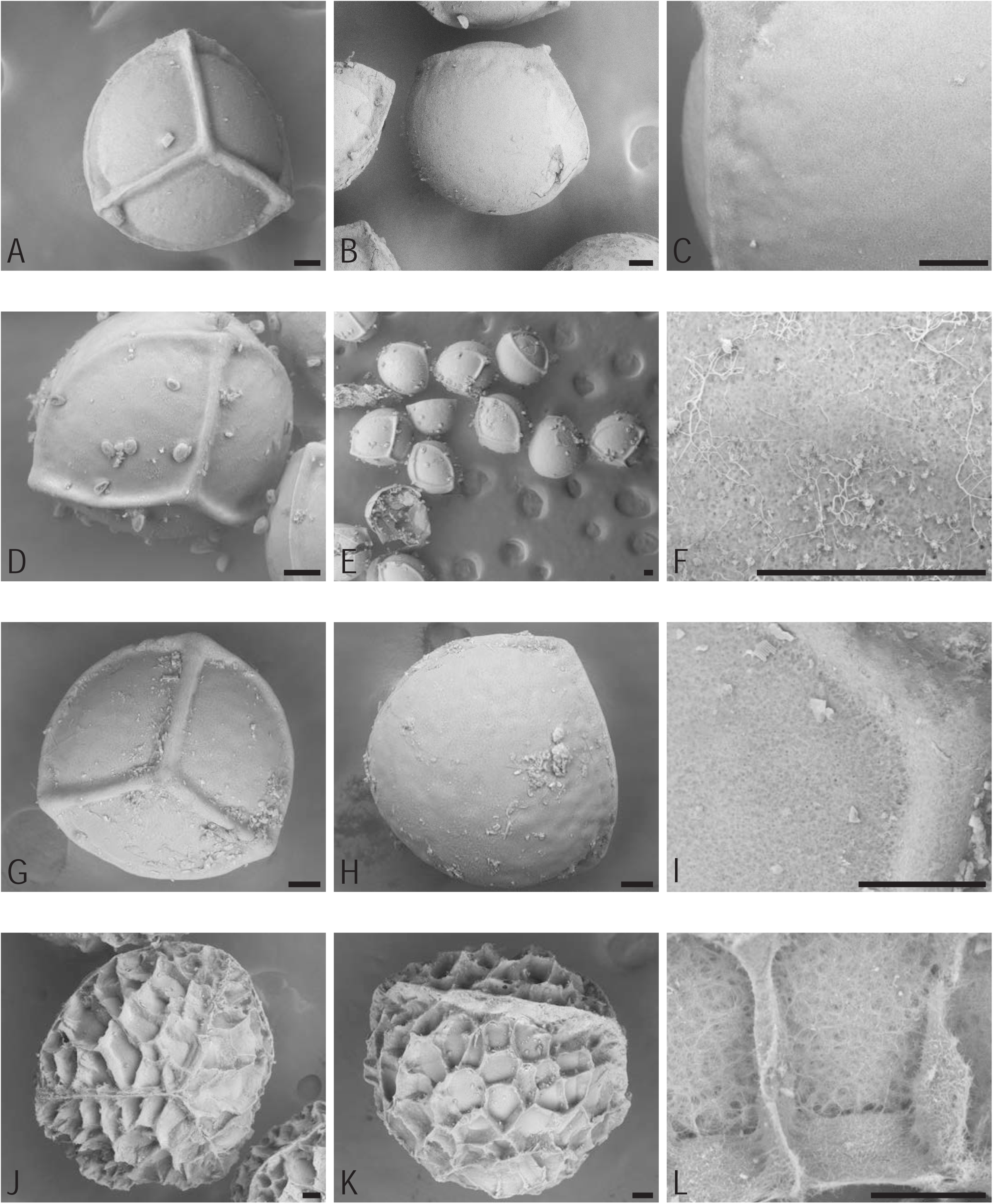
Species of clade E. All scale bars are 50 μm. A-C *Isoetes lechleri* ELS42, A proximal side, B distal side, C magnified surface structure on distal side. D-F *Isoetes herzogii* ELS62, D proximal side, E several spores, some showing distal side, F magnified surface structure on proximal side, with fungal hyphae. G-I *Isoetes boliviensis* ELS37, G proximal side, H distal side, I magnified surface structure and proximal ridges. J-L *Isoetes killipii* ELS90, J proximal side, K distal side, L magnified surface structure on proximal side.

#### *Isoetes herzogii* U.Weber ELS62 (Fig. 19D-F)

340-380 µm. Proximal ridges medium tall, medium wide and forming points where they meet the equatorial ridge. Equatorial ridge medium, not very distinct. Ornamentation is levigate, on both sides devoid of features. No obvious girdle. Surface structure is dense with pores.

#### *Isoetes boliviensis* U.Weber ELS37 (Fig. 19G-I)

380-390 µm. Proximal ridges wide, tall and forming points when meeting the equatorial ridge. Equatorial ridge of medium height and even. Ornamentation is levigate, on both sides devoid of features. No obvious girdle. Surface structure is dense with pores.

#### *Isoetes killipii* C.V.Morton ELS90 (Fig. 19J-L)

660-740 µm. Proximal ridges narrow, tall, sharp and blending into the ornamentation. Equatorial ridge narrow, tall and sharp. Ornamentation is reticulate. On both sides paper-thin walls of uneven height. No obvious girdle. Surface structure is a network of fibers with short loose ones.

#### *Isoetes engelmannii* A.Braun ELS19 (Fig. 20A-C)

460-490 µm. Proximal ridges very thin, tall and sharp with small protrusions along them. Equatorial ridge very thin, tall, sharp and blending into the ornamentation. Ornamentation is reticulate. On both sides paper-thin walls of very uneven height with frayed ends. No obvious girdle. Surface structure is dense with a network of thin fibers as outmost layer.

**Figure 20.**
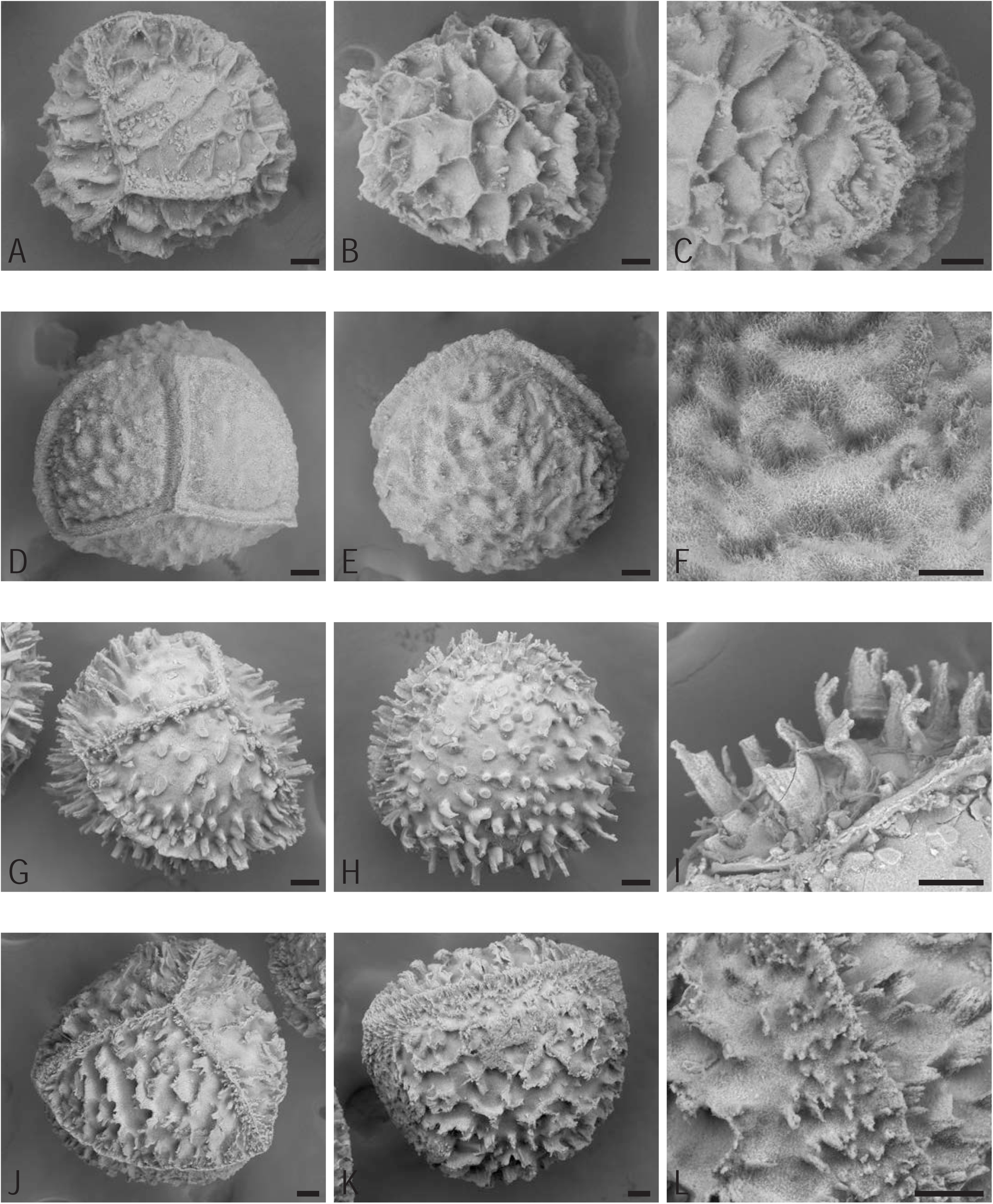
Species of clade E. All scale bars are 50 μm. A-C *Isoetes engelmannii* ELS19, A proximal side, B distal side, C magnified ornamentation on distal side. D-F *Isoetes virginica* ELS124, D proximal side, E distal side, F magnified surface structure on proximal side, proximal ridge above and equatorial ridge to the left. G-I *Isoetes echinospora* ELS13, G proximal side, H distal side, I magnified surface structure on proximal side. J-L *Isoetes lacustris* ELS123, J proximal side, K distal side, L microspores just below the equatorial ridge.

#### *Isoetes virginica* N.Pheiff. ELS124 (Fig. 20D-F)

460-520 µm. Proximal ridges wide-ish, quite low, with a crowning ridge, no ornamentation on them and pulled into points where they meet the equatorial ridge. Equatorial ridge quite tall and lumpy. Ornamentation is rugulate. On proximal side tuberculate and covered in fur. On distal side rugulate (but hard to see the exact ridge forms because it is very hairy). Larger distal side than proximal side. Girdle with no ornamentation. Surface structure is hairy.

#### *Isoetes echinospora* Durieu ELS13 (Fig. 20G-I)

430-460 µm. Proximal ridges very thin, tall, with a crowning ridge and unevenly spaced protrusions. Equatorial ridge very thin, sharp, tall and uneven. Ornamentation is echinate. On both sides protrusions are often joining into broader bands rather than solitary needles and they tend to fork and occasionally bend near the tip. No obvious girdle. Surface structure is dense with a network of thin fibers as outmost layer.

#### *Isoetes lacustris* L. ELS123 (Fig. 20J-L)

600-630 µm. Proximal ridges very tall and narrow with small protrusions crowding the sides. Equatorial ridge very tall, narrow, sharp, ribbed and uneven. Ornamentation is rugulate, on both sides very tall with thin and frayed walls. Girdle with densely packed echinate protrusions. Surface structure is a dense network of fibers with short loose ones.

#### *Isoetes azorica* Durieu ex Milde ELS89 (Fig. 21A-C)

410-480 µm. Proximal ridges wide and low, occasionally with short protrusions. Equatorial ridge regular and quite smooth. Ornamentation is rugulate, on proximal side with low, narrow and uneven ridges. On distal side the ridges are wider and sometimes almost giving a reticulate impression. Girdle of sparse, low, cristate structures. Surface structure is a dense network of fibers with evenly spaced short loose ones.

**Figure 21.**
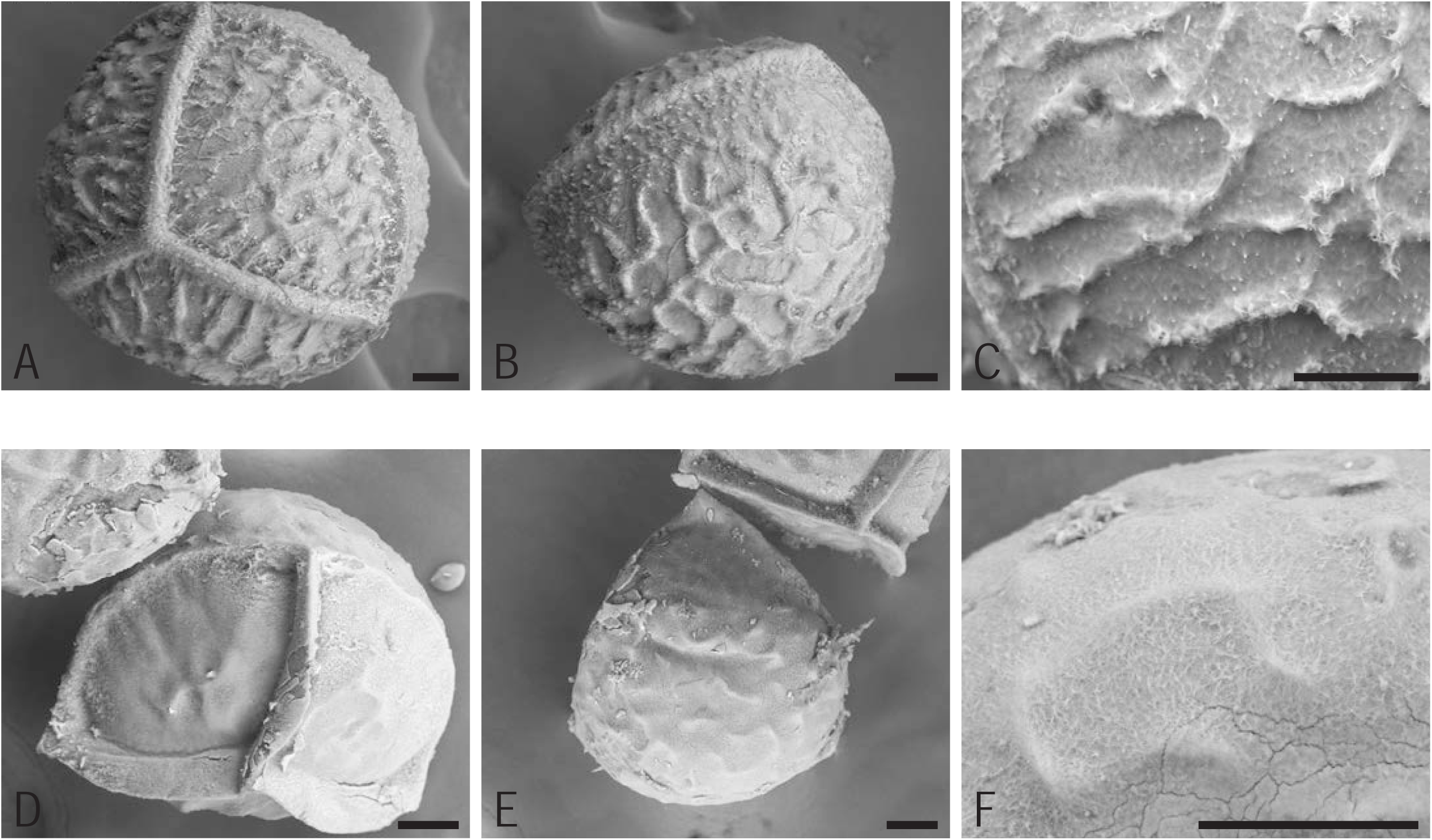
Species of clade E. All scale bars are 50 μm. A-C *Isoetes azorica* ELS89, A proximal side, B distal side, C magnified ornamentation on distal side. D-F *Isoetes melanospora* ELS56, D proximal side, E distal side, F magnified surface structure on proximal side, proximal ridge above and equatorial ridge to the left.

#### *Isoetes melanospora* Engelm. ELS56 (Fig. 21D-F)

310-350 µm. Proximal ridges thin, exceedingly tall and forming points when meeting the equatorial ridge. Equatorial ridge quite thin and not very distinct. Ornamentation is rugulate, on proximal side almost levigate except for some ridges that are more like creases on the surface. On distal side rugulate with very low and short ridges. No obvious girdle. Surface structure is dense with a network of thin fibers as outmost layer.

### Phylogenetic analysis

The results of the phylogenetic analysis (Fig. 2) are congruent with those of Larsén *et al*. (2022), which is wholly expected since the dataset used here is a subselection of the same sequences. Seven sequences were, however, newly produced for the present study (denoted with a star * in Supporting Information Table S1) in order to, as much as possible, complement information on spore morphology with phylogenetic information based on molecular data.

The South African species *Isoetes wormaldii* is sister to the remaining genus (bootstrap value BS=100). Clade A sensu Larsén and Rydin (2016) is sister to remaining species (except *I. wormaldii*) (BS=100). Within clade A (BS=100), the South African *I. capensis* is sister to remaining species (BS=100). Remaining species in clade A (BS=100) form two sister clades, of which one comprises a set of South and Central American species (BS=100). Its sister (BS=100) comprises species from tropical (to southern) Africa, India, tropical Asia and Australia. Results in the South-Central American clade are not all well supported, whereas those of the latter clade are for the most part well-supported.

In clade B sensu Larsén and Rydin (2016) (BS=100), the Mediterranean species *I. durieui* and *I. histrix* (BS=100) are sister to the remaining species (BS=100). The North American west coast species *I. nuttallii* and *I. orcuttii* (BS=100) are sister to remaining species (BS=100), which includes species from southern (to tropical) Africa, Madagascar, India and the Mediterranean region. Relationships within this part of clade B are unclear as support values are generally low.

The Italian endemic *I. malinverniana*, clade C sensu Larsén and Rydin (2016), is sister to remaining species (BS=100), which comprise clades D and E sensu Larsén and Rydin (2016).

Clade D sensu Larsén and Rydin (2016) (BS=100) is divided into two subclades, one comprising species from Australia, New Zealand and India (BS=96), and the other of species from eastern and tropical Asia (BS=100), within which one of the included species from New Guinea is sister to the remaining species (BS=88).

Within clade E sensu Larsén and Rydin (2016) (BS=100), two species from the northern Andes, *I. andicola* and *I. andina*, (BS=100) are sister to a clade (BS=99) that comprises species from South America, Central America, North America, and the circumboreal region. Relationships in this latter clade are mostly poorly supported.

## DISCUSSION

### General findings

The main goal of the present study was to document the variation of megaspore morphology in Isoetaceae and evaluate its potential relevance for understanding relationships and evolution in the lineage through time. And crucially, our study is done with feet on more solid ground regarding phylogeny compared to previous work on the entire family; most sampled specimens have been sequenced and their placement in the phylogeny is known, meaning that no matter the species identification, the sample’s position in the tree is known. Although this was not possible for some very old plant material, effort was made to assess the phylogenetic position also of samples for which we lack molecular data. While there are general features common to all megaspores of the family, there is in addition a surprising amount of morphological variation. Some of this variation appears clade-specific and possible to explain in a macroevolutionary perspective (descent with modification); other features may instead be related for example to the ecology of the species, intraspecific variation, and hybridisation/introgression/polyploidisation.

In contrast with the monolete microspores, the megaspores of all species of *Isoetes* are trilete, which is in agreement with previous observations (among many Pfeiffer 1922, Musselman 2002). As found here, the megaspores of *Isoetes* usually range in size between 400 and 600 µm with an average size difference within a sample of about 50 µm, but for example the species of two early diverging clades in clade B (*I. orcuttii, I. nuttallii, I. olympica, I. longissima, I*. sp. ELS116) have smaller megaspores, down to 300 µm. By contrast, one of the species in the sister clade of the remaining clade B (*I. durieui*) has among the largest megaspores we detected, up to 710 µm. Megaspores up to 750 µm, and 780 µm also occur in *I. malinverniana* (sister to clades D+E) and a few species in clade E. It is often argued that polyploidy is related to increased cell size (e.g., spore size) but in a synopsis of karyological data in *Isoetes,* Troìa (2001) points out that there are contradictory results in the literature regarding the correlation between megaspore size and ploidy level in *Isoetes* (referring to Kott and Britton 1983, Watanabe *et al*. 1996). Ploidy levels may often vary within species of *Isoetes* (Troìa 2001, Larsén and Rydin 2016, Troìa *et al*. 2016) and we have no karyological data for our samples but as assessed based on literature information, our detected interspecific differences in spore size may well be related to chromosome number. Several of our investigated species with the largest spores (e.g., *I durieui, I. malinverniana, I. novo-granadensis*) are reported to have high chromosome numbers: 2n=44, 55, 110; 2n=44; and 2n=126-132, respectively (Troìa *et al*. 2016). The opposite is true for the species mentioned above as those with the smallest spores in our study; they are all reported to have 2n=22 (Troìa *et al*. 2016). These indications are in line with results in Pereira *et al*. (2015) who found significant positive correlation between spore size and ploidy level for Brazilian species of *Isoetes*.

Differences among species of *Isoetes* regarding the ornamentation and surface texture/structure of the megaspore are considerable. The South African narrow endemic *Isoetes wormaldii*, recently shown to be sister to the remaining *Isoetes* (Larsén *et al*. 2022, Wikström *et al*. 2023) have a unique foveolate megaspore ornamentation (Fig. 3A-C) not seen or reported for any other species of *Isoetes*. This goes in concert with previous conclusions on this species, e.g., its (for *Isoetes*) unusual leaf morphology (Larsén *et al*. 2022) and the fact that it is so molecularly divergent from the remaining genus that adding it to phylogenetic analyses approximately doubled the average median node depth to the *Isoetes* crown group (Wikström *et al*. 2023), a result that is roughly reproduced also in the present study (inset phylogram in Fig. 2).

### Descriptive spore morphology

The terms we have used for description of megaspore ornamentation mainly follow Hickey (1986b), though not in every respect. We recognize nine of Hickey’s (1986b) 12 terms (baculate, cristate, echinate, levigate, pustulate, retate, reticulate, rugulate, and tuberculate) for megaspore ornamentation, and in addition three more (foveolate, spiculate and umbellate). A pustulate ornamentation is here assessed as the most common megaspore ornamentation in *Isoetes*; it occurs in most species of clade B, is common in clade A and is present in some species in the C-D-E clade (Fig. 2). This contradicts conclusions in early work (Pfeiffer 1922) where the majority of the species are defined as having tuberculate megaspore ornamentation. Hickey (1986) pointed out that the four categories of megaspore ornamentation used by Pfeiffer (1922) do not cover the diversity of the genus. He made, for example, a distinction between tuberculate (straight inclined sides with an acute apex) and pustulate (convex sides with an obtuse, rounded apex) ornamentation (Hickey 1986b). As assessed here, actual tuberculate spore ornamentation occurs primarily in clade A (Fig. 2).

Baculate ornamentation is mainly present in South American species of clade A; echinate and levigate ornamentation occur mainly in clade E (rarely in clade D); retate ornamentation in a few species of clades B and D; rugulate ornamentation is only present in clades D and E; cristate and reticulate ornamentation are rare and occur only in a few species each. Foveolate ornamentation (in *Isoetes wormaldii*) is to our knowledge a new term for describing megaspore ornamentation in *Isoetes*.; its spores have previously been described as reticulate (Pfeiffer 1922, Duthie 1929), but no other spores in the genus looks like those of *Isoetes wormaldii*. Pfeiffer (1922) describes the megaspores of *I. japonica* as having “large foveolate markings”, and choice of terminology can always be discussed, but comparing our spores of *I. japonica* (Fig. 13A-C) with those of *I. wormaldii* (Fig. 3A-C), clearly shows the distinct difference in appearance between the two. “Spiculate” is perhaps self-explanatory. One could argue that echinate would mean the same as spiculate, but the spores historically described as echinate have projections without distinct apexes, more similar to columns than spicules. Distinct spicules occur in the North American sister-species *I. orcuttii* and *I. nuttallii* of clade B. The ornamentation form umbellate was described for *Isoetes* spores by Singh *et al*. (Singh *et al*. 2021a, Singh *et al*. 2021b), and is here assessed for two African samples of clade B.

The terminology that has been used to describe the surface texture/structure of megaspores of *Isoetes* has often varied between studies; no general standard has been established. One exception is the texture described as cobweb, which was first mentioned by Wanntorp (1970), later also used by Marsden (1979), and again in the present study. The feature occurs in the megaspores of *Isoetes wormaldii* and all non-South American species of clade A (Fig. 2). Over the remaining phylogeny, a hairy surface texture is common in clade B and in some species of clades D-E. A network-like texture occurs mostly in clades D-E but also in some species of the other clades. A further of seven different kinds of surface textures/structures are identified here but most of them are rare, occurring only in one to a few species or small clades (Fig. 2).

### Evolutionary trends and spore morphology of the different clades

Despite some potential clade-defining characteristics, the variation in megaspore ornamentation and surface texture/structure in *Isoetes* often appears unclear from a geographical and macroevolutionary perspective (Fig. 2), and it seems unlikely that a reconstruction analysis could have defined the ancestral states of features with confidence. One type of ornamentation keeps reappearing over the phylogeny of *Isoetes* (pustulate ornamentation) and the cobwebby type of surface texture/structure is shared between *I. wormaldii* and all non-South American species of clade A (and reappears also in clade C) (Fig. 2). A tentative speculation for the ancestral states of these characters in *Isoetes* could thus be a pustulate megaspore ornamentation with a cobwebby surface structure. However, the sister to the rest of the genus (*Isoetes wormaldii*) does not have pustulate megaspore ornamentation but a unique foveolate pattern, and further studies and analyses are clearly needed. It was beyond the scope of the present study to compare our results with those of its living sister genus *Selaginella*, but relevant work on *Selaginella* is available (e.g. Korall and Taylor 2006) and for future work, such comparison may be an important component for an improved understanding of the evolution of megaspore morphology in the isoetalean clade.

#### Clade A — “the Gondwana clade”

The very specific cobweb-like structure that is the dominating surface texture in clade A was described by Wanntorp (1970) in the first scanning electron microscope studies of megaspores of *Isoetes*. He argues that south-western African species of *Isoetes* either, as in *I. kersii*, have the “cobwebby” surface texture, or as in *I. giessii* and *I. alstonii* a denser network of strands that form “short acute spines” (Wanntorp 1970). These descriptions are entirely congruent with our findings, where *I. kersii* (currently considered a synonym of *I*. *schweinfurthii*), represented by sample EL035=ELS93, has a cobwebby surface texture and falls in clade A, whereas *I. giessii* and *I. alstonii* with different surface textures belong in clade B. Apart from the subclade with South American species, all sampled species of clade A have the cobwebby spore texture and tuberculate or pustulate ornamentation.

Duthie (1929) described the megaspore ornamentation of *Isoetes capensis* and other species (*I. stellenbossiensis* and *I. stephansenii*) that are more recently shown to belong in the *I. capensis* clade (e.g. Hoot *et al*. 2006, Larsén and Rydin 2016) as reticulate. We consider the ornamentation of *I. capensis* (Fig. 3D-F) to be best described as pustulate and images in Duthie (1929) do not contradict that. The remaining African, Indian and Australian specimens in clade A included here all have similar ornamentation, either pustulate or tuberculate and with a cobwebby surface structure. One of our two samples of *I. coromandelina* (ELS72) has been determined to subspecies (*I. coromandelina* subsp. *macrotuberculata*), whereas the other (ELS117) is undetermined to subspecies (Figs 2, 5A-F). The former sample is collected in Australia and the latter on mainland Asia (Thailand), which means that the former should be correctly determined to *I. coromandelina* subsp. *macrotuberculata* while the latter should represent *I. coromandelina* subsp. *coromandelina*. However, based on our results on spore morphology and comparison with documentation in the literature (Marsden 1979, Singh *et al*. 2021b) the determination to subspecies of these samples, and/or the distinct geographical division of these two subspecies as currently circumscribed (i.e. *I. coromandelina* subsp. *macrotuberculata* present in northern Australia; *I. coromandelina* subsp. *coromandelina* in mainland Asia), could be questioned. Additional studies of *Isoetes coromandelina* with a substantially increased sampling from its entire geographic distribution is needed to clarify the matter.

Within the South American subclade of clade A, most spores have a baculate ornamentation and a surface texture described as network, but there is variation among species and some have a unique combination of features not seen in any other species of *Isoetes*. The two sister species *Isoetes clavata* ELS121 (Fig. 3J-L) and *Isoetes pedersenii* ELS104 (Fig. 4A-C) exemplifies this; the former has baculate ornamentation and a surface texture of curly fibers whereas the latter has tuberculate ornamentation and a hairy surface texture. A comprehensive understanding of phylogeny and evolution of spore morphology in South American *Isoetes* requires additional studies though, not least since South American species of *Isoetes* fall into two distantly related clades, clades A and E (Fig. 2 and previous work, e.g. Rydin and Wikström 2002, Larsén and Rydin 2016, Pereira *et al*. 2017a, Larsén *et al*. 2022). Further, ample recent work has shown that the diversity in South American *Isoetes* is substantial and at least some of the newly described species (see e.g. Pereira *et al*. 2016, Pereira *et al*. 2019) clearly belong in clade A (Pereira *et al*. 2017a).

#### Clade B — “the cosmopolitan clade”

In the two first clades that separate away from the rest of Clade B (Fig. 2) the ornamentation is either pustulate (*I. histrix*), retate (*I. durieui*) or spiculate (*I. orcuttii* and I*. nuttallii*), and the surface texture/structure is sponge-like, covered in thick thorns or bearing a remarkable resembles to melted cheese. Several of these features are unique to these two subclades of clade B, but a pustulate ornamentation is common in many clades and a retate ornamentation is in addition found in one species of clade D (*I. stevensii*). Interestingly, our sample of *I. orcuttii* could potentially be a hybrid specimen; it had two types of spores in the same sporangium: levigate spores and spores that were densely spiculate. In the third clade to diverge in clade B (comprising *I. olympica*, *I. longissima* and a sample undetermined to species), megaspore morphology is the same as in most species of clade B: a pustulate ornamentation and a hairy surface texture. Only *I. olympica* has a surface texture that is more of a network with loose fibers, rather than hairy.

Ample work has investigated the spore morphology and dispersal of the species in these clades and provides valuable and detailed information (see e.g. Hickey 1986b, Troìa and Raimondo 2009, Troìa *et al*. 2012, Greuter and Troìa 2015, Troìa 2016, Troìa and Rouhan 2018). As often in *Isoetes*, a complicating factor when assessing evolutionary patterns and processes in Mediterranean and North American species is that they may belong in different major clades, in this case either in clade B or in clade E (e.g. Hoot *et al*. 2004, Larsén and Rydin 2016, Pereira *et al*. 2017a, Troìa *et al*. 2019, Larsén *et al*. 2022). However, since most of the specimens in the remaining clade B, both African and European species, have a pustulate ornamentation and a hairy surface structure, our tentative hypothesis is that the former is an ancestral feature in *Isoetes* and the latter a synapomorphy of clade B (although certainly not unique to clade B). A few exceptions exist: baculate ornamentation in the Indian species *Isoetes dixitii*, umbellate ornamentation in two African specimens and a surface structure that is not hairy but rather consisting of a network of fibers in two (distantly related) African specimens (*I. alstonii* and *I. natalensis*). All these surface textures/structures are strikingly different from the cobwebby nature common in megaspores produced by species of clade A, and the difference could probably be used as a rough estimate of clade affinity of African *Isoetes*, perhaps even species affinity.

The term “umbellate” was first used in *Isoetes* by Singh *et al*. (2021a, 2021b) to describe the megaspore ornamentation of *I. dixitii*; however, our investigated spores of this species clearly have a baculate ornamentation. Instead, we find an umbellate ornamentation in the spores of two of our African samples representing *I. aequinoctialis* and *I. welwitschii*, respectively. Although these samples both belong in clade B, they do not form a clade. It could also be mentioned that our interpretations on megaspore morphology appear entirely congruent with photos and written information in the works by Duthie (1929) and Wanntorp (1970), and mostly also Pfeiffer (1922) although the quality of the photos in the latter study often prevents precise comparisons. Additional studies are required to assess diversity and variation in megaspore ornamentation and surface structure in clade B, including whether intraspecific variation exists. Furthermore, an overall revision of diversity and species delimitation in African *Isoetes* is needed (Larsén *et al*. 2022). The species *Isoetes schweinfurthii* and *I. welwitschii* are clearly not monophyletic as currently circumscribed; sampled specimens may be resolved in clade A or clade B (Fig. 2 and Larsén *et al*. 2022). And as assessed from the results of the present study, the same may hold for additional species, for example *Isoetes aequinoctialis* (Fig. 2).

#### Clade C — “the malinverniana clade”

In clade C, here only represented by *Isoetes malinverniana*, we find the only species outside of clade A that has cobweb as surface structure on its megaspores, but what is more unique and interesting with this species is the dimorphic megaspore ornamentation. There are examples within *Isoetes* of the ornamentation varying between the proximal and the distal side of the spore, but in *Isoetes malinverniana* the two forms of ornamentation both occur on the proximal side. There are three low pustules closest to the proximal conjunction, each in their own radial area. The rest of the spore is covered in bacules that even on occasion blend into the equatorial ridge. Further, this species has the unusual characteristic of a raised layer of netted rough fibers that partly covers its ornamentation. Spores from several specimens of *I. malinverniana* were studied to make sure that this characteristic is representative for the species and not an abnormal state. The second species that belongs in clade C, *Isoetes anatolica* (Bolin *et al*. 2008, Larsén and Rydin 2016) was not included in the present study but previous work has indicated several similarities between *I. malinverniana* and *I. anatolica* regarding megaspore morphology (Prada and Rolleri 2005), although the extra, raised layer of fibers *I. malinverniana* (Fig. 12J-L) may to some extent disguise the similarity.

#### Clade D — “the Australasian clade”

Clades D and E contain an increased amount of diversity of ornamentation types and surface structures compared to clades A-C. Clade D is divided into two well-supported subclades, and the members of these groups show differences regarding megaspore morphology. The megaspores of the clade with representatives from Australia, New Zealand and India (Fig. 2, lower subclade of clade D) have either pustulate or rugulate ornamentation and their surface structure is hairy or occasionally a network of fibers. The diversity of megaspore morphology is even greater in the clade with representatives from New Guinea, Philippines, Japan and China (Fig. 2, upper subclade of clade D); its megaspores can have a retate, reticulate, rugulate, or echinate ornamentation. The diversity of ornamentation types in this latter clade is, however, not matched by the surface structure as they all have a network of fibers.

One feature appears to be a synapomorphy for the upper subclade of clade D, and that is an upward-directed equatorial ridge on the megaspores. The generality of this feature in this subclade is also evident from Marsden’s comprehensive thesis work (1979) on the morphology and taxonomy of *Isoetes* in Australasia (see e.g. figure 209 of *I. neoguineensis* Baker and figure 153 of *I. philippinensis* in Marsden, 1979), and a comparison of our documentation of megaspore ornamentation with his indicates full agreement.

#### Clade E — “the American clade”

The first clade to diverge from the remaining species of clade E includes *Isoetes andicola* that shows an ornamentation type unique to some species of clade E: a levigate megaspore ornamentation (Fig. 15G-I). Apparently, a levigate megaspore ornamentation typically appears in association with a dense surface structure with occasional pores (Fig. 2). *Isoetes andicola* was once placed in its own genus because of a perceived general distinctness from the remaining Isoetaceae (Amstutz 1957). A levigate megaspore ornamentation of *I. andicola* is however not unique to this species; it occurs in several other species of clade E and, based on literature information, probably also in some species of clade D. The megaspores of *Isoetes hopei*, placed in clade D in previous work (Larsén *et al*. 2022), were not included in the present study but are described as “almost smooth” and as “ornamentation … lacking” by Marsden (1979) and Croft (1980). The same holds for *I. gunnii*, phylogenetically placed in clade D (Larsén *et al*. 2022) and with a megaspore ornamentation described by Hickey as levigate (Hickey 1986b).

The functional consequences of loss of megaspore ornamentation appear mostly negative, for example by making it less likely that microspores stick to the megaspores and are transported together (see e.g. Fig. 7L), which is otherwise common (Musselman 2002, Troìa 2016), and perhaps also by decreasing the probability that the megaspores stick to, and are transported by, any dispersal agent. In the extinct arborescent isoetaleans, loss of megaspore ornamentation is coupled with the reduction of the number of megaspores per sporangium to one, i.e. a cease of megaspores as dispersal units and the evolution of a “seed-like” dispersal system (Bateman *et al*. 1992). Such systems are, however, not present in extant *Isoetes*, and the advantage (if any) of smooth (i.e. levigate) spores in extant *Isoetes* must be another. Dispersal of the entire megasporangium as a unit, as well as dispersal via the digestive tract of animals, are reported for *Isoetes* (see e.g. Troìa 2016 for a summary of dispersal in *Isoetes*), and may conceivably make spore ornamentation superfluous, but the potential correlation with such dispersal strategies and levigate megaspores is not known, at least not to us.

The sister to the levigate *I. andicola* (*I. andina*) has another unique appearance of its megaspores: a cristate ornamentation with a hairy surface structure (Fig. 15J-L). The cristate condition was originally described as tubercles or spines that are somewhat extended into ridges (Pfeiffer 1922) but *I. andina* is mentioned by Hickey (1986b) as one of the key examples of a cristate ornamentation.

It is premature to discuss evolutionary patterns in the remaining clade E since the phylogeny has proven difficult to resolve, perhaps due to hybridization and polyploidy (see Larsén *et al*. 2022 for a recent discussion on the phylogeny of the clade). Furthermore, clade E clearly includes ample variation in megaspore morphology; pustulate, reticulate, rugulate, levigate and echinate ornamentations occur and surface structures are either hairy, dense, dense with pores or a network of fibers (Figs 2, 16-21). This variation is also evident from ample studies in the literature (among many, Pfeiffer 1922, Kott and Britton 1983, Hickey 1985, 1986b, Taylor 1993, Musselman and Knepper 1994, Macluf *et al*. 2003, Troìa and Raimondo 2009, Pereira *et al*. 2012, Troìa *et al*. 2012, Pereira and Labiak 2013, Brunton 2015, Greuter and Troìa 2015, Pereira *et al*. 2017b, Troìa and Rouhan 2018, Troìa *et al*. 2019). To fully grasp phylogeny and spore evolution in clade E requires ample additional work on a substantially increased sample of taxa, preferably in combination with genomic data from the same samples, which would permit investigations of intergenomic discord and reasons thereof, such as ILS and hybridization/introgression (e.g. Rieseberg and Soltis 1991, Stull *et al*. 2020, Thureborn *et al*. 2022, Thureborn *et al*. 2024).

### Relationships with extinct members of the isoetalean (rhizomorphic) clade

In a genus such as *Isoetes* with seemingly little morphological variation (and putatively extensive parallelism) it can be very difficult to more specifically relate the living species to fossil taxa (Pigg 1992, 2001). Moreover, the isoetalean lineage (i.e. the rhizomorphic lycopsids) and a majority of its historical diversity is extremely old, at least 350 million years, possibly up to almost 400 million years as assessed from analyses of paleobotanical as well as molecular data (e.g. Bateman *et al*. 1992, Pigg 1992, Larsén and Rydin 2016, DiMichele *et al*. 2022), and this temporal distance will complicate homology assessments. Attempts to compare the morphology of extant and extinct forms are (rightfully?) at risk of being questioned simply because of the age gap.

Unfortunately, it has also been shown that the feature in focus in the present study, spore morphology, is less phylogenetically informative than other reproductive features (i.e. cone morphology) in the isoetalean lineage, because spore features may often be autapomorphic, homoplasious, and/or replacing each other entirely rather than successively accumulating change (Bateman *et al*. 1992), which complicates understanding of evolution. Based on our literature survey we further find that even though dispersed spores, like pollen, are widespread in fossil assemblages across the world, only fossil spores that have been discovered in situ in *Isoetes*-like plants are of value for understanding the macroevolutionary history of the isoetalean clade since the uncertainties of taxonomic affinity are too great for dispersed spores.

Despite these obstacles, we find the fossil megaspores of *Isoetes reticulata* Hill, discovered from late Oligocene to early Miocene deposits in Tasmania (Hill 1988) of particular interest. Hill (1988) described them as having “*a strongly developed triradiate ridge, surface pattern reticulate. Reticulations larger and more prominent on distal spore face. Megaspore surface consisting of a dense layer of mesh-like filaments*.”, and compared them with megaspores of extant *Isoetes* species occurring in Australia and Asia today (as described by Marsden 1979). Hill (1988) found no clear similarities beyond basic ornamentation type to any extant species. However, when studying spore ornamentation of extant species across the phylogeny, we find some clear cases of similarity to the fossil *I. reticulata*. Like the extant Australasian species of the upper subclade of clade D (Fig. 2), *I. reticulata* has an equatorial ridge that is tilted upwards. And the reticulate megaspore ornamentation of the fossil is similar to that of the extant *I. neoguineensis* (e.g. figure 209 in Marsden, 1979) and *I. japonica* (Fig. 13A-C); the ornamentation pattern is more similar to *I. neoguineensis* while the structure of the walls building the ornamentation more closely resembles those of *I. japonica*. In addition, the equatorial ridge of the fossil is sinuous and “lumpy”, as it is in *I neoguineensis*.

While the fossil megaspores of *I. reticulata* is not an exact match of those of any extant species (indeed that would have been most surprising), we argue based on optimization of these characters on the phylogeny that the fossil *I. reticulata* belongs to clade D, possibly as sister to the subclade that includes *I. neoguineensis* and *I. japonica*. While we find it interesting to note that accepting this argumentation would yield a node age of clade D of about 20-25 Ma, i.e. within the estimated confidence interval of 20–50 Ma for clade D in Larsén and Rydin (2016), later work in our lab (Wikström *et al*. 2023) has shown that node age estimation in *Isoetes* is a notoriously difficult scientific problem. Finding justifications for a calibration point to absolute time within the crown group in analyses of divergence times of clades may offer a solution.

## SUPPORTING INFORMATION

Supporting information (Table S1) is available at *Botanical Journal of the Linnean Society* online.

## AUTHOR CONTRIBUTIONS

EL and CR designed the work. EL conducted data curation, SEM studies and phylogenetic analyses. AK produced the molecular data. All authors interpreted the results. EL wrote the first draft of the manuscript. All authors commented and edited the text.

## DATA AVAILABILITY

All molecular data used in the present study are deposited at GenBank. Accessions are given in the Supporting Information Table S1.

## ACKNOWLEDGEMENTS

We thank Dr Kjell Jansson (the Electron Microscopy Centre EMC, Stockholm University) for technical assistance, and Dr Sylvain Razafimandimbison (The Swedish Museum of Natural History) and the herbaria BM, BR, C, L, MEL, P, S, W, WU for access to plant material.

