## Supplemental Table 1 for "Spore morphology and evolution in *Isoetes* (Isoetales)"

**Supporting Information Table S1.** List of taxon names, distributions, DNA voucher information (including area and year of collection), lab identity numbers, and accession numbers for sequences used in the analysis. References are given for sequences taken from GenBank. Taxon names and authority follow Troïa *et al*. [1] and Tropicos [2] with a few exceptions [see nomenclatural notes in reference 3]. Distributions of *Isoetes* were extracted from Troïa *et al*. [1], Tropicos [2], GBIF.org [4], and SANBI's Red List of South African Plants [5]. Area names follow the World Geographical Scheme for Recording Plant Distributions [6]. An exception is the Mediterranean distribution, which refers to an occurrence in either one of the 22 sovereign countries in Europe, Africa and Temperate Asia that borders the Mediterranean Sea.

| DNA Id | Spore Id | Taxon | Distribution | DNA Voucher | Area and year of collection | *ndhC-ndhK* | *rbcL* | *rpoC1* | *ycf1* | *ycf66* | $trnV^{UAC}$ | nrITS |
|---|---|---|---|---|---|---|---|---|---|---|---|---|
| EL002 | ELS035 | *Isoetes aequinoctialis* Welw. ex A.Braun | Southern and Trop. Africa | Kornas 3453 (BR) | Zambia 1973 | OM691715 | OM691983 | OM870154 | OM870000 | OM691793 | OM691883 | OM719899 |
| EL051 | ELS054 | *Isoetes aequinoctialis* Welw. ex A.Braun | Southern and Trop. Africa | Giess 15279 (S) | Nambia 1978 | OM691751 | OM692024 | OM870196, OM870292, OM870348 | OM870042 | OM691828 | OM691920 | OM719939 |
| EL165 | ELS004 | *Isoetes aequinoctialis* Welw. ex A.Braun | Southern and Trop. Africa | Hall 907 (BM) | Ghana 1965 | - | - | - | - | - | - | OP437695* |
| EL121 | ELS100 | *Isoetes alpina* Kirk | New Zealand | Melville & Connor 6250 (L) | New Zealand 1962 | - | - | - | - | - | - | KT288369 |
| EL114 | ELS106 | *Isoetes alstonii* C.F.Reed & Verdc. | Southern and Trop. Africa | Milne-Redhead & Taylor 10003 (BR) | Tanzania 1956 | - | - | New* | - | - | - | - |
| EL059 | - | *Isoetes andicola* (Amstutz) L.D.Gómez | Western South America | Marshall s.n. (BM) | Peru 1961 | - | OM692031 | OM870203, OM870299, OM870355 | OM870049, OM870122 | OM691835 | OM691927 | OM719945 |
| - | ELS108 | *Isoetes andicola* (Amstutz) L.D.Gómez | Western South America | Hutchinson & Tovar 4244 (C) | Peru 1964 | - | - | - | - | - | - | - |
| EL118 | ELS103 | *Isoetes andina* Spruce ex Hook. | Western & Northern South America | Larsen & Eriksen 19 (WU) | Ecuador 1985 | OM691785 | OM692079 | OM870251, OM870382 | OM870097, OM870142 | OM691875 | OM691971 | OM719983 |
| - | ELS024 | *Isoetes andina* Spruce ex Hook. | Western & Northern South America | Cuatrecasas & Jeramillo 25977 (BM) | Colombia 1961 | - | - | - | - | - | - | - |
| EL026 | ELS089 | *Isoetes azorica* Durieu ex Milde | The Azores | Gonalves 2611 (BM) | The Azores 1971 | OM691731 | OM692004 | OM870175 | OM870020 | OM691810 | OM691899 | OM719919 |
| EL030 | ELS046 | *Isoetes biafrana* Alston | Trop. Africa | Le Testu 3016 (BM) | C. Afr. Rep. 1951 | OM691735 | OM692008 | OM870179 | OM870024 | OM691814 | OM691903 | OM719923 |
| EL098 | ELS118 | *Isoetes bolanderi* Engelm. | Western Canada, Northwestern & Southwestern USA | Munz s.n. (C) | USA 1926 | OM691775 | OM692064 | OM870235, OM870375 | OM870081 | OM691863 | OM691956 | OM719972 |
| EL011 | ELS037 | *Isoetes boliviensis* U.Weber | Western South America | Hickey 753 (W) | Bolivia 1980 | - | OM691990 | OM870161, OM870277, OM870334 | OM870006, OM870111 | - | - | OM719905 |
| EL038 | ELS096 | *Isoetes capensis* A.V.Duthie | Southern Africa | Duthie s.n. (BM) | South Africa 1934 | - | - | - | OM870031 | - | - | OM719929 |
| EL095 | ELS121 | *Isoetes clavata* U.Weber | Northern South America | Boudrie 3922 (P) | French Guiana 2003 | OM691774 | OM692061 | OM870233, OM870325, OM870374 | OM870079 | OM691862 | OM691954 | OM719971 |
| EL099 | ELS117 | *Isoetes coromandelina* L.f. | Indian Subcontinent, Indo-China, Australia | Larsen 8398 (C) | Thailand 1961 | - | OM692065 | OM870236, OM870326 | OM870082, OM870137 | OM691864 | OM691957 | - |
| - | ELS006 | *Isoetes coromandelina* L.f. | Indian Subcontinent, Indo-China, Australia | Wallace 5915 (BM) | Myanmar 1945 | - | - | - | - | - | - | - |
| - | ELS016 | *Isoetes coromandelina* L.f. | Indian Subcontinent, Indo-China, Australia | Goswami s.n. (BM) | India (unknown | - | - | - | - | - | - | - |

| DNA Id | Spore Id | Taxon | Distribution | DNA Voucher | Area and year of collection | *ndhC-ndhK* | *rbcL* | *rpoC1* | *ycf1* | *ycf66* | *trnV*[UAC] | nrITS |
|---|---|---|---|---|---|---|---|---|---|---|---|---|
| | | | | | year) | | | | | | | |
| EL135 | - | *Isoetes coromandelina* L.f. subsp. *coromandelina* | India | Srivastava 450268 (S) | India 2000 | - | OM692089 | OM870261, OM870332 | OM870107 | - | - | OM719993 |
| EL086 | ELS072 | *Isoetes coromandelina* L.f. subsp. *macro-tuberculata* C.R.Marsden | Australia (northern) | Walsh & Coles 4441 (MEL) | Australia 1996 | OM691771 | OM692056 | OM870228, OM870320, OM870371 | OM870073, OM870133 | OM691856 | OM691949 | OM719966 |
| EL136 | ELS102 | *Isoetes dixitii* Shende | India | Patil 6 (K) | India 1973 | - | OM692090 | OM870262 | OM870108, OM870145 | - | OM691981 | OM719994 |
| EL022 | - | *Isoetes drummondii* A.Braun | Australia | Beauglehole 75018 (MEL) | Australia 1983 | OM691728 | OM692000 | OM870171 | OM870016 | OM691807 | OM691896 | OM719915 |
| - | ELS027 | *Isoetes drummondii* A.Braun | Australia | Chinnock P1036 (BM) | Australia 1974 | - | - | - | - | - | - | - |
| EL027 | ELS044 | *Isoetes durieui* Bory | Mediterranean | Byfield s.n. (BM) | Turkey 1992 | OM691732 | OM692005 | OM870176 | OM870021 | OM691811 | OM691900 | OM719920 |
| EL028 | ELS045 | *Isoetes durieui* Bory | Mediterranean | De Retz 65141 (BM) | France 1972 | OM691733 | OM692006 | OM870177 | OM870022 | OM691812 | OM691901 | OM719921 |
| - | ELS015 | *Isoetes durieui* Bory | Mediterranean | Humphries & Richardson 103 (BM) | Italy 1973 | - | - | - | - | - | - | - |
| EL013 | - | *Isoetes echinospora* Durieu | Europe, Northern America, Asia-Temperate | Ford 609 (W) | Canada 2006 | OM691721 | OM691992 | OM870163 | OM870008 | OM691799 | OM691889 | OM719907 |
| - | ELS013 | *Isoetes echinospora* Durieu | Europe, Northern America, Asia-Temperate | Villaret 17232 (BM) | Germany 1955 | - | - | - | - | - | - | - |
| GB | - | *Isoetes engelmannii* A.Braun | Northcentral, Northeastern & Southeastern USA | Taylor s.n. (MIL) | USA 1995 | - | - | - | - | - | - | DQ479978 |
| - | ELS019 | *Isoetes engelmannii* A.Braun | Northcentral, Northeastern & Southeastern USA | Kezer s.n. (BM) | USA 1937 | - | - | - | - | - | - | - |
| EL113 | ELS107 | *Isoetes flaccida* var. *chapmanii* Engelm. | Southeastern USA | Godfrey 61963 (BR) | USA 1962 | - | OM692076 | OM870247 | OM870093, OM870140 | OM691872 | OM691967 | OM719981 |
| EL101 | ELS115 | *Isoetes gardneriana* Kunze ex A.Braun | Brazil, Southern South America | Pedersen 19654 (C) | Argentina 1984 | OM691777 | OM692067 | OM870238, OM870327 | OM870084 | OM691866 | OM691959 | OM719974 |
| EL044 | ELS050 | *Isoetes giessi* Launert | Namibia | Giess, Volk & Bleissner 5564 (S) | Nambia 1963 | OM691745 | OM692018 | OM870190, OM870286, OM870343 | OM870036 | OM691823 | OM691914 | OM719934 |
| EL005 | ELS036 | *Isoetes giessii* Launert | Namibia | Giess, Volk & Bleissner 5564 (BR) | Namibia 1963 | OM691716 | OM691985 | OM870156 | OM870001 | OM691794 | OM691884 | OM719901 |
| EL074 | ELS062 | *Isoetes herzogii* U.Weber | Bolivia | Halls s.n. (BM) | Bolivia 1984 | OM691763 | OM692045 | OM870217, OM870310, OM870364 | OM870062 | OM691847 | OM691940 | OM719958 |
| - | ELS014 | *Isoetes histrix* Bory | Mediterranean | Reverchon 275 (BM) | Greece 1884 | - | - | - | - | - | - | - |
| EL168 | ELS005 | *Isoetes histrix* Bory | Mediterranean | Reverchon 3699 (BM) | France 1894 | | - | - | - | - | - | OP437697* |
| EL080 | ELS077 | *Isoetes howellii* Engelm. | Northwestern & Southwestern USA & Western Canada | Thorne & Lathrop 37942 (BM) | USA 1969 | OM691768 | OM692051 | OM870223, OM870270, OM870316 | OM870068, OM870129 | - | - | - |

| DNA Id | Spore Id | Taxon | Distribution | DNA Voucher | Area and year of collection | *ndhC-ndhK* | *rbcL* | *rpoC1* | *ycf1* | *ycf66* | *trnV*$^{UAC}$ | nrITS |
|---|---|---|---|---|---|---|---|---|---|---|---|---|
| EL058 | ELS069 | *Isoetes humilior* F.Muell. ex A.Braun | Australia | Darbyshire 134 (BM) | Australia 1961 | OM691755 | OM692030 | OM870202, OM870266, OM870298, OM870354 | OM870048, OM870121 | OM691834 | OM691926 | OM719944 |
| EL103 | ELS114 | *Isoetes japonica* A.Braun | Eastern Asia | Amano 328 (C) | Japan 1986 | OM691779 | OM692069 | OM870240, OM870329, OM870377 | OM870086 | OM691867 | OM691961 | OM720001, OM719889 |
| EL024 | ELS090 | *Isoetes killipii* C.V.Morton | Northern & Western South America | Grubb & Guymer P. 37 (BM) | Colombia 1957 | - | OM692002 | OM870173, OM870282, OM870339 | OM870018 | - | OM691897 | OM719917 |
| EL081 | ELS078 | *Isoetes kirkii* A.Braun | New Zealand | Chinnock P447 (BM) | New Zealand 1972 | OM691769 | OM692052 | OM870224, OM870271, OM870317, OM870369 | OM870069, OM870130 | OM691853 | OM691946 | OM719963 |
| EL064 | ELS123 | *Isoetes lacustris* L. | Europe, Northern America, Asia-Temperate | Taylor 4904 (BM) | USA 1983 | OM691759 | OM692036 | OM870208, OM870304, OM870359 | OM870054 | OM691840 | OM691932 | OM719950 |
| EL025 | ELS042 | *Isoetes lechleri* Mett. | Western South America | Fernandez-Casas & Molero 6619 (BM) | Bolivia 1982 | OM691730 | OM692003 | OM870174 | OM870019 | OM691809 | OM691898 | OM719918 |
| EL052 | ELS017 | *Isoetes longissima* Bory subsp. *longissima* | Mediterranean | De Retz 65107 (BR) | France 1972 | OM691752 | OM692025 | OM870197, OM870293, OM870349 | OM870043 | OM691829 | OM691921 | OM719940 |
| EL032 | - | *Isoetes malinverniana* Ces. & De Not. | Italy | Raynal 20885 (BR) | Italy 1978 | OM691736 | OM692009 | OM870180 | OM870025 | OM691815 | OM691904 | OM719924 |
| | ELS011 | *Isoetes malinverniana* Ces. & De Not. | Italy | Mattivolo 1606 (BM) | Italy 1910 | - | - | - | - | - | - | - |
| | ELS126 | *Isoetes malinverniana* Ces. & De Not. | Italy | Gola & Ferrari 5599 (W) | Italy 1910 | - | - | - | - | - | - | - |
| | ELS127 | *Isoetes malinverniana* Ces. & De Not. | Italy | Gola & Ferrari 5599 (W) | Italy 1910 | - | - | - | - | - | - | - |
| EL122 | ELS099 | *Isoetes melanopoda* J.Gay & Durieu | Northcentral, Northeastern & Southeastern USA | Thomas 88386 (L) | USA 1984 | OM691787 | OM692081 | OM870253, OM870384 | OM870099 | - | OM691973 | OM719985 |
| EL056 | ELS056 | *Isoetes melanospora* Engelm. | Southeastern USA | Spongberg & Boufford 1726 (BM) | USA 1982 | OM691754 | OM692028 | OM870200, OM870296, OM870352 | OM870046 | OM691832 | OM691924 | OM719942 |
| EL093 | - | *Isoetes melanotheca* Alston | West Tropical Africa | Raynal 7693 (P) | Senegal 1961 | - | - | - | OM870077 | OM691860 | - | OM719887 |
| - | ELS003 | *Isoetes melanotheca* Alston | West Tropical Africa | Pitot s.n. (BM) | Guinea 1950 | - | - | - | - | - | - | - |
| EL061 | ELS084 | *Isoetes mexicana* Underw. | Mexico | Pringle 8796 (BM) | Mexico 1904 | - | OM692033 | OM870205, OM870301 | OM870051, OM870123, OM870149 | OM691837 | OM691929 | OM719947 |
| EL060 | ELS083 | *Isoetes montezumae* A.A.Eaton | Mexico | McVaugh 13650 (BM) | Mexico 1952 | OM691756 | OM692032 | OM870204, OM870300, OM870356 | OM870050 | OM691836 | OM691928 | OM719946 |
| EL115 | ELS105 | *Isoetes muelleri* A.Braun | Australia | Stajsic 908 (MEL) | Australia 1993 | - | OM692077 | OM870248, OM870331 | OM870094, OM870141 | OM691873 | OM691968 | - |
| EL082 | - | *Isoetes natalensis* Baker | Madagascar, Southern Africa | Burrows 3734 (BM) | Namibia 1987 | - | OM692053 | OM870225, OM870272 | OM870070, OM870131 | - | OM691947 | OM719964 |
| - | ELS113 | *Isoetes natalensis* Baker | Madagascar, Southern | Perrier de la Bathie 13648 | Madagascar | - | - | - | - | - | - | - |

| DNA Id | Spore Id | Taxon | Distribution | DNA Voucher | Area and year of collection | *ndhC-ndhK* | *rbcL* | *rpoC1* | *ycf1* | *ycf66* | *trnV*[UAC] | nrITS |
|---|---|---|---|---|---|---|---|---|---|---|---|---|
| | | | Africa | (C) | 1921 | | | | | | | |
| EL020 | - | *Isoetes neoguineensis* Baker | Papua New Guinea | Craven 2717 (MEL) | Papua New Guinea 1974 | OM691726 | OM691998 | OM870169 | OM870014 | OM691805 | OM691894 | OM719913 |
| L5 | - | *Isoetes novo-granadensis* H.P.Fuchs | Northern & Western South America | Holm-Nielsen, L. B. et al. 5470 (L) | Ecuador 1973 | - | - | - | - | - | - | KT288385 |
| - | ELS031 | *Isoetes novo-granadensis* H.P.Fuchs | Northern & Western South America | Santa & Escobar 1113 (BM) | Colombia 1985 | - | - | - | - | - | - | - |
| EL049 | ELS053 | *Isoetes nuttallii* A.Braun ex Engelm. | Western Canada, Northwestern & Southwestern USA | Macoun 86378 (S) | Canada 1908 | OM691749 | OM692022 | OM870194, OM870290 | OM870040, OM870118 | - | OM691918 | OM719997, OM719883 |
| EL046 | - | *Isoetes olympica* A.Braun | Syria, Turkey | Samuelsson 4566 (S) | Syria 1933 | OM691747 | OM692020 | OM870192, OM870288, OM870345 | OM870038, OM870117 | OM691825 | OM691916 | OM719936 |
| - | ELS088 | *Isoetes olympica* A.Braun | Syria, Turkey | Nydegger 11519 (BM) | Turkey 1976 | - | - | - | - | - | - | - |
| EL045 | ELS051 | *Isoetes orcuttii* A.A.Eaton | Southwestern USA, Mexico | Carter 471 (S) | USA 1934 | OM691746 | OM692019 | OM870191, OM870287, OM870344 | OM870037 | OM691824 | OM691915 | OM719935 |
| EL016 | - | *Isoetes palmeri* H.P.Fuchs | Northern & Western South America | Small 155 (W) | Colombia 1993 | - | OM691995 | OM870166, OM870279, OM870336 | OM870011, OM870112 | OM691802 | OM691892 | OM719910 |
| - | ELS032 | *Isoetes palmeri* H.P.Fuchs | Northern & Western South America | Jermy 17483 (BM) | Colombia 1986 | - | - | - | - | - | - | - |
| EL105 | ELS112 | *Isoetes panamensis* Maxon & C.V.Morton | Central America, Western South America, Brazil | Irwin, Harley & Smith 31615 (C) | Brazil 1971 | - | OM692070 | OM870241, OM870378 | OM870087, OM870139, OM870152 | OM691868 | OM691962 | OM719976 |
| EL106 | ELS111 | *Isoetes paraguayensis* H.P.Fuchs (nom. nud.) | --- | Pedersen 7624 (C) | Paraguay 1965 | OM691780 | OM692071 | OM870242 | OM870088 | OM691869 | OM691963 | OM719977 |
| EL117 | ELS104 | *Isoetes pedersenii* H.P.Fuchs ex E.I.Meza & Macluf | Western & Southern South America, Brazil | Abbott 16374 (WU) | Bolivia 1995 | OM691784 | OM692078 | OM870250, OM870381 | OM870096 | OM691874 | OM691970 | OM719982 |
| EL036 | ELS094 | *Isoetes philippinensis* Merr. & L.M.Perry | Malesia | Price 500 (BM) | Philippines 1969 | OM691739 | OM692012 | OM870184 | OM870029 | OM691818 | OM691908 | OM719927 |
| EL034 | ELS092 | *Isoetes pitotii* Alston | West Tropical Africa | Hall 3696 (BM) | Ghana 1967 | - | - | OM870182 | OM870027, OM870114 | - | OM691906 | OM719882 |
| EL109 | ELS110 | *Isoetes rhodesiana* Alston | East Tropical Africa | Vollesen 4586 (C) | Tanzania 1977 | OM691782 | OM692073 | OM870244, OM870380 | OM870090 | OM691870 | OM691965 | - |
| EL065 | ELS085 | *Isoetes riparia* Engelm. ex A.Braun | Eastern Canada, Northeastern & Southeastern USA | Jermy 12487 (BM) | Canada 1975 | - | OM692037 | OM870209, OM870360 | OM870055 | - | OM691933 | OM719951 |
| EL085 | ELS073 | *Isoetes saccharata* Engelm. | Northeastern USA | Eaton s.n. (MEL) | USA 1903 | - | - | - | - | - | - | New* |
| EL039 | ELS049 | *Isoetes sampathkumaranii* L.N.Rao | Indian Subcontinent | Goswami s.n. (BM) | India (unknown year) | OM691741 | OM692014 | OM870186 | OM870032 | OM691820 | OM691910 | OM719930 |
| EL076 | ELS075 | *Isoetes schweinfurthii* A.Braun | Southern and Trop. Africa | Gilbert 878 (BM) | Ethiopia 1975 | OM691764 | OM692047 | OM870219, OM870312, OM870365 | OM870064 | OM691849 | OM691942 | OM719959 |

| DNA Id | Spore Id | Taxon | Distribution | DNA Voucher | Area and year of collection | *ndhC-ndhK* | *rbcL* | *rpoC1* | *ycf1* | *ycf66* | *trnV*$^{UAC}$ | nrITS |
|---|---|---|---|---|---|---|---|---|---|---|---|---|
| EL077 | ELS063 | *Isoetes schweinfurthii* A.Braun | Southern and Trop. Africa | Garrod & Sanusi 6309 (BM) | Nigeria 1977 | OM691765 | OM692048 | OM870220, OM870313, OM870366 | OM870065, OM870128 | OM691850 | OM691943 | OM719960 |
| EL035 | ELS093 | *Isoetes schweinfurthii* A.Braun. | Southern and Trop. Africa | Kers 3130 (BM) | Namibia 1968 | OM691738 | OM692011 | OM870183 | OM870028 | OM691817 | OM691907 | OM719926 |
| EL163 | ELS001 | *Isoetes schweinfurthii* A.Braun. | Southern and Trop. Africa | Drummond & Rutherford 7557 (BM) | Zambia 1961 | - | - | - | - | - | - | OP437693* |
| EL164 | ELS002 | *Isoetes schweinfurthii* A.Braun. | Southern and Trop. Africa | Kornas 6272 (BM) | Nigeria 1977 | - | - | - | - | - | - | OP437694* |
| - | ELS029 | *Isoetes schweinfurthii* A.Braun. | Southern and Trop. Africa | Giess 7615 (BM) | Namibia 1963 | - | - | - | - | - | - | - |
| EL111 | ELS1009 | *Isoetes sinensis* T.C.Palmer | China | [Illegible] s.n. (C) | China 1927 | - | OM692075 | OM870246, OM870330 | OM870092 | - | - | OM719980 |
| EL048 | ELS052 | *Isoetes* sp. | | Suksdorf s.n. (S) | USA 1909 | OM691748 | OM692021 | OM870193, OM870263, OM870289, OM870346 | OM870039 | OM691826 | OM691917 | OM719937 |
| EL054 | ELS055 | *Isoetes* sp. | Europe, Northern America, Asia-Temperate | Øllgaard & Pedersen 180 (BM) | Denmark 1954 | - | OM692026 | OM870198, OM870294, OM870350 | OM870044, OM870119 | OM691830 | OM691922 | OM719941 |
| EL097 | ELS119 | *Isoetes* sp. | | Rajaonary, Ravololomanana & Porembski 205 (S) | Madacascar 2015 | - | OM692063 | OM870234 | OM870080, OM870136, OM870151 | - | OM691955 | - |
| EL100 | ELS116 | *Isoetes* sp. | Western Mediterranean | Ortiz & Pueche 1469 (C) | Spain 1977 | OM691776 | OM692066 | OM870237, OM870376 | OM870083 | OM691865 | OM691958 | OM719973 |
| EL124 | ELS098 | *Isoetes stevensii* J.R.Croft | Papua New Guinea | Schodde 1843 (C) | Papua New Guinea 1961 | - | OM692083 | OM870255, OM870385 | OM870101, OM870143 | OM691877 | OM691975 | OM719987 |
| EL062 | ELS124 | *Isoetes virginica* N.Pfeiff. | South-central & Southeastern USA | Spongberg 1743 (BM) | USA 1982 | OM691757 | OM692034 | OM870206, OM870302, OM870357 | OM870052 | OM691838 | OM691930 | OM719948 |
| EL029 | ELS087 | *Isoetes welwitschii* A.Braun | Southern and Trop. Africa | Wingfield 2032 (BM) | Tanzania 1972 | OM691734 | OM692007 | OM870178 | OM870023 | OM691813 | OM691902 | OM719922 |
| EL128 | ELS067 | *Isoetes welwitschii* A.Braun | Southern and Trop. Africa | Razafimandimbison, Razafindrahaja, Atalahy & Swenson 2142 (S) | Madagascar 2018 | OM691790 | OM692087 | OM870259 | OM870105 | OM691881 | OM691979 | OM719991 |
| EL129 | ELS068 | *Isoetes welwitschii* A.Braun | Southern and Trop. Africa | Razafimandimbison, Razafindrahaja, Atalahy & Swenson 2151 (S) | Madagascar 2018 | OM691791 | OM692088 | OM870260 | OM870106 | OM691882 | OM691980 | OM719992 |
| EL167 | ELS033 | *Isoetes welwitschii* A.Braun | Southern and Trop. Africa | Gilbert & Thulin 904 (BM) | Ethiopia 1975 | - | - | - | - | - | - | OP437696* |
| EL057 | ELS081 | *Isoetes wormaldii* Sim. | South Africa (Eastern Cape) | Pocock 20009 (BM) | South Africa 1955 | - | OM692029 | OM870201, OM870265, OM870297, OM870353 | OM870047 | OM691833 | OM691925 | - |
| GB | | *Dendrolycopodium obscurum* (L.) A.Haines | --- | --- | --- | MH549637[7] | MH549637[7] | MH549637[7] | MH549637[7] | MH549637[7] | MH549637[7] | - |
| GB | | *Diphasiastrum digitatum* (Dill. ex | --- | --- | --- | MH549638[7] | MH549638[7] | MH549638[7] | MH549638[7] | MH549638[7] | MH549638[7] | - |

| DNA Id | Spore Id | Taxon | Distribution | DNA Voucher | Area and year of collection | *ndhC-ndhK* | *rbcL* | *rpoC1* | *ycf1* | *ycf66* | *trnV*[UAC] | nrITS |
|---|---|---|---|---|---|---|---|---|---|---|---|---|
| | | A.Braun) Holub | | | | | | | | | | |
| GB | | *Huperzia lucidula* (Michx.) Trevis. | --- | --- | --- | AY660566[8] | AY660566[8] | AY660566[8] | AY660566[8] | AY660566[8] | AY660566[8] | – |
| GB | | *Huperzia serrata* (Thunb.) Trevis. | --- | --- | --- | KX426071[9] | KX426071[9] | KX426071[9] | KX426071[9] | KX426071[9] | KX426071[9] | – |
| GB | | *Lycopodium clavatum* L. | --- | --- | --- | MH549642[7] | MH549642[7] | MH549642[7] | MH549642[7] | MH549642[7] | MH549642[7] | - |
| GB | | *Selaginella bisulcata* Spring | --- | --- | --- | MH598531[10] | MH598531[10] | MH598531[10] | MH598531[10] | - | - | - |
| GB | | *Selaginella doederleinii* Hieron. | --- | --- | --- | MH598532[10] | MH598532[10] | MH598532[10] | MH598532[10] | - | - | - |
| GB | | *Selaginella hainanensis* X.C.Zhang & Noot. | --- | --- | --- | - | MH598533[10] | MH598533[10] | MH598533[10] | - | - | - |
| GB | | *Selaginella indica* (Milde) R.M.Tryon | --- | --- | --- | - | MK156801[11] | MK156801[11] | MK156801[11] | - | - | - |
| GB | | *Selaginella kraussiana* (Kunze) A.Braun | --- | --- | --- | MH549643[7] | MH549643[7] | MH549643[7] | MH549643[7] | - | - | - |
| GB | | *Selaginella lepidophylla* (Hook. & Grev.) Spring | --- | --- | --- | - | MK089531[7] | MK089531[7] | MK089531[7] | - | - | - |
| GB | | *Selaginella lyallii* (Hook. & Grev.) Spring | --- | --- | --- | - | MK156800[10] | MK156800[10] | MK156800[10] | - | - | - |
| GB | | *Selaginella moellendorffii* Hieron. | --- | --- | --- | MG272484[10] | MG272484[10] | MG272484[10] | MG272484[10] | - | - | - |
| GB | | *Selaginella pennata* (D.Don) Spring | --- | --- | --- | MH598534[10] | MH598534[10] | MH598534[10] | MH598534[10] | - | - | - |
| GB | | *Selaginella remotifolia* Spring | --- | --- | --- | MH598535[10] | MH598535[10] | MH598535[10] | MH598535[10] | - | - | - |
| GB | | *Selaginella sanguinolenta* (L.) Spring | --- | --- | --- | - | MH598536[10] | MH598536[10] | MH598536[10] | - | - | - |
| GB | | *Selaginella tamariscina* (P.Beauv.) Spring | --- | --- | --- | - | MH598537[10] | MH598537[10] | MH598537[10] | - | - | - |
| GB | | *Selaginella uncinata* (Desv.) Spring | --- | --- | --- | MG272483[10] | MG272483[10] | MG272483[10] | MG272483[10] | - | - | - |
| GB | | *Selaginella vardei* H.Lév. | --- | --- | --- | - | MG272482[11] | MG272482[11] | MG272482[11] | - | - | - |

**Notes**: * (asterisk) denotes sequences newly produced for the present study. Accessions that begin with OM were produced by Larsén *et al.* [3].
